# The role of insulators in transgene transvection in *Drosophila*

**DOI:** 10.1101/565846

**Authors:** Pawel Piwko, Ilektra Vitsaki, Ioannis Livadaras, Christos Delidakis

## Abstract

Transvection is the phenomenon where a transcriptional enhancer activates a promoter located on the homologous chromosome. It has been amply documented in *Drosophila* where homologues are closely paired in most, if not all, somatic nuclei, but it has been known to rarely occur in mammals as well. We have taken advantage of site-directed transgenesis to insert reporter constructs into the same genetic locus in *Drosophila* and have evaluated their ability to engage in transvection by testing many heterozygous combinations. We find that transvection requires the presence of an insulator element on both homologues. Homotypic *trans*-interactions between four different insulators can support transvection: the *gypsy insulator* (*GI*), *Wari, Fab-8* and *1A2*; *GI* and *Fab-8* are more effective than *Wari* or *1A2*. We show that in the presence of insulators, transvection displays the characteristics that have been previously described: it requires homologue pairing, but can happen at any of several loci in the genome; a solitary enhancer confronted with an enhancerless reporter is sufficient to drive transcription; it is weaker than the action of the same enhancer-promoter pair in *cis* and it is further suppressed by *cis*-promoter competition. Though necessary, the presence of homotypic insulators is not sufficient for transvection; their position, number and orientation matters. A single GI adjacent to both enhancer and promoter is the optimal configuration. The addition of a heterologous insulator in one homolog can positively or negatively influence transvection strength. The local landscape of enhancers and promoters is also important, indicative of complex insulator-enhancer-promoter interactions.

## Introduction

Transgenesis and reporter genes are crucial tools for studying gene regulation. However, a transgene is often susceptible to interactions with surrounding chromatin leading to significant variations in expression patterns and levels, depending on the site of insertion in the genome (Levis *et al.* 1985). In order to factor-out these “position effects” multiple lines of transgenic animals must be studied. More recently, this problem has been tackled by two strategies now widely used in *Drosophila*. One relies on the ability to guide integration of a transgene to a specific location in the genome via the use of the *ΦC31*-mediated integration system (Thorpe and Smith 1998; Groth *et al.* 2004; Pfeiffer *et al.* 2008; Kvon 2015; Markstein *et al.* 2008). By targeting all reporters to the same “landing site”, they are directly comparable and only a single transgenic line per construct needs to be analysed. The second way to minimize genomic position effects utilizes insulator sequences in transgenesis vectors. Insulator DNA elements were first identified by their potential to block gene function when interposed between enhancer and promoter (Kellum and Schedl 1992; Cai and Levine 1995; Kuhn *et al.* 2003) and to insulate transgenes from effects of surrounding chromatin (Kellum and Schedl 1991; Roseman *et al.* 1993, 1995). These faculties of insulators are linked to their strong propensity for interaction with each other (Kuhn *et al.* 2003; Chetverina *et al.* 2008; Kyrchanova *et al.* 2008a, 2011). Such interactions are proposed to form chromatin loops bridging even distant loci at the base of the loop and isolating interactions inside the loop from interactions outside the loop (Byrd and Corces 2003; Blanton *et al.* 2003; Doyle *et al.* 2014; Cubeñas-Potts and Corces 2015).

We started using both of these approaches to characterize enhancer modules of two neighboring *Drosophila* genes *E*(*spl*)*m7* and *E*(*spl*)*m8*. To that end we generated a series of reporter constructs flanked by two copies of the insulator sequence from the *gypsy* transposon (*gypsy* insulator, GI) and integrated each construct into the same *attP* locus. When we tested two different reporters in a heterozygous configuration, we were surprised to find they were able to affect each other’s expression in *trans*. The ability of enhancers to activate promoters in the homologous locus is called transvection and was first reported in *Drosophila*, as a phenomenon of pairing-dependent intragenic (unexpected) complementation in loci like *Ultrabithorax* (*Ubx*) (Lewis 1954; Martínez-Laborda *et al.* 1992), *decapentaplegic* (*dpp*) (Gelbart 1982), *yellow* (Geyer *et al.* 1990) and *white* (Babu and Bhat 1980). Subsequent studies employing randomly integrated P-element transgenes showed that the *Drosophila* genome is generally permissive to enhancer action in *trans* (Chen *et al.* 2002; Kravchenko *et al.* 2005) offering a first glimpse of the molecular basis of this phenomenon and reconfirming the need for somatic homologue pairing (or synapsis). Homologue synapsis is not limited to germline meiotic cells in *Drosophila* (and dipterans in general), but is very common in somatic tissues. This is in contrast to mammals, where somatic homologue synapsis seems to happen only in special occasions and accordingly only sporadic cases of transvection have been reported (McKee 2004; Heride *et al.* 2010; Apte and Meller 2012; Stratigi *et al.* 2015; Joyce *et al.* 2016). The development of site-specific integration methods in *Drosophila* (such as the *ΦC31*-based recombination method) has recently rekindled the interest in transvection (Lee and Wu 2006; Bateman *et al.* 2012a; Mellert and Truman 2012; Fujioka *et al.* 2016). However, the mechanism of transvection is still poorly understood.

Using our series of reporter transgenes, we decided to search for sequence determinants of transvection. We found that this interaction is dependent on the presence of homotypic insulator DNA elements on both homologues. Our transgenesis vectors contained two such insulators: the Gypsy Insulator (GI) commonly utilized to protect transgenes from genomic position effects, and the Wari Insulator (WI) carried in the 3’ part of the mini-*white* marker gene. Two other insulators were also found to support transvection, while insulator removal from either of the *trans*-interacting transgenes abrogated transvection at five discrete genomic loci. While necessary, the presence of insulators was not sufficient to produce a robust transvection outcome. Parameters like the number, position and orientation of GIs relative to the *trans*-interacting enhancers and promoters proved to be of paramount importance. The implications of these results on the design of transgenes and on the broader role of insulators in transcription will be discussed.

## Materials and Methods

### Plasmid constructs

See attached **Supplementary Materials and Methods** file.

### Fly maintenance and stocks

Flies were maintained under standard conditions at 25°C. Stocks containing attP docking sites used for the integration of *attB* plasmids: *attP40* (RRID:*BDSC_25709*), *attP2* (RRID:*BDSC_25710*) (Groth *et al.* 2004; Markstein *et al.* 2008), *VK2* (RRID:*BDSC_9723*), *VK13* (RRID:*BDSC_24864*), *VK37* (RRID:*BDSC_24872*), *VK40* (RRID:*BDSC_35568*) (Venken *et al.* 2006); each carrying, and if not - crossed to, a chromosome expressing *ΦC31* integrase under the control of *nanos* derived from *attP40* stock (RRID:*BDSC_25709*) (Bischof *et al.* 2007). The *su*(*Hw*) mutant effects were assayed in the animals transheterozygous for *su*(*Hw*)^*e04061*^ null allele (RRID:<u>BDSC_18224</u>) (Thibault *et al.* 2004) and *su*(*Hw*)^*2*^ strong hypomorphic allele resulting in the ten times decreased *su*(*Hw*) expression (RRID:<u>BDSC_983</u>) (Parkhurst *et al.* 1988; Georgiev *et al.* 1997).

### Integration of attB plasmids into attP fly lines

All plasmids in this study were integrated into a unique attP landing site specified in the text and figure legends for each transgene. Microinjection was performed as previously described (Ringrose 2009). A solution of 500 ng/μl plasmid DNA was microinjected into *nanos- ΦC31; attP* fly embryos. Flies that grew to adulthood were crossed with *y w* flies. Depending on the injected DNA construct, the mini-*white* or *3xP3-dsRed* marker was used for subsequent screening and tracking the transgene.

### Immunostaining and microscopy

Fixation and immunohistochemistry of embryos and larval tissues was performed according to standard protocols. CNSs and imaginal disks were dissected from late third instar larvae, fixed with 4% paraformaldehyde and labeled with rabbit polyclonal anti-GFP (Minotech Biotechnology) and mouse anti-β-galactosidase (Promega Cat# Z3781, RRID:<u>AB_430877</u>) primary antibodies. Goat anti-rabbit IgG secondary antibody, Alexa488-conjugated (Thermo Fisher Scientific Cat# A-11034, RRID:<u>AB_2576217</u>) was used for GFP detection. Goat anti-mouse, Alexa633-conjugated (Thermo Fisher Scientific Cat# A-21052, RRID:<u>AB_2535719</u>) or donkey anti-mouse, Alexa647-conjugated (Thermo Fisher Scientific Cat# A-31571, RRID:<u>AB_162542</u>) secondary antibodies were used for β-galactosidase detection. Samples were imaged on a Leica SP8 confocal platform using a 20x Oil Immersion Objective with fixed zoom levels for each tissue type (CNS, wing and eye disks). The images were pseudo-colored in green (GFP), red (LacZ) and blue (DsRed). All samples within each figure were fixed and immunostained at the same time. Scanning of all figure samples was performed using identical microscope and software settings and, when possible, completed within one imaging session to enable semi-quantitative comparison. Where scanning of all figure samples within one session was not possible, replicates of the samples from two chosen genotypes were re-scanned together with the remaining samples in the next scanning session ensuring that the replicated samples are comparable. Images were manipulated using ImageJ (pseudocoloring, rotation and maximum intensity projection z-stacks) and arranged into data sets using Adobe Photoshop CC 2017 and Microsoft PowerPoint. Note that we used two different z-projections for some wing disk images. For the top part, containing the wing pouch, a full z-projection of the sample was done, while the bottom part, containing the notum and hinge, encompassed only sections containing the adult muscle precursors (AMPs). This was done to avoid confusing AMP expression with expression in the overlying tegula, a sensory organ primordium. Whereas AMPs and tegula can be easily distinguished in the 3D confocal stacks, they merge to one cluster upon z-projection. Since enhancer e7 activity is specific for the AMPs and not the tegula, we excluded the tegula sections from the z-projections shown.

### Luciferase assays

Luciferase activity was measured using the Promega Luciferase Assay System Kit (Cat# E153A). CNSs and imaginal disks from ten late third instar larvae were collected in 200 μl of 1X lysis reagent CCLR for each sample. Samples were collected over a series of days and stored at −80°C until five independent samples were collected for each genotype. Samples were defrosted, put on ice and homogenized using Kontes pestles. Homogenized samples were incubated at room temperature for 10 minutes and then centrifuged for 5 minutes to pellet tissue remains. Obtained homogenates were subsequently measured for luciferase activity and total protein content. 20 μl of each homogenized sample was mixed with 50 μl of Promega Luciferase Assay Reagent and promptly measured on single tube luminometer (Turner Designs TD-20/20). Total protein was measured using the Pierce BCA Protein Assay Kit (Cat# 23225). 10 μl of each homogenized sample was mixed with 200 μl BCA Working Reagent on clear-bottomed 96-well plates (Costar) and incubated at 37°C for 1 hr. The plates were transferred at room temperature for 10 min to allow the reactions to stabilize. Absorbance was measured on the Awareness Technology ChroMate Microplate Reader at 562 nm. Three replica plates were averaged for each sample. A standard curve was produced with BSA dilutions in Promega 1X lysis reagent CCLR. Luciferase activity from each sample was normalized to its total protein content.

## Data Availability

All plasmids and fly strains are available upon request. The authors affirm that all data necessary for confirming the conclusions of the article are present within the article, figures, and tables.

## Results

### Enhancer element analysis of E(spl)m7 and E(spl)m8 reveals a transvection phenomenon

This work began with the aim to dissect the transcriptional regulation of two adjacent genes in the *E*(*spl*) complex, *E*(*spl*)*m7* and *E*(*spl*)*m8*, both encoding transcription factors involved in many aspects of the response to Notch signaling. These are small, intronless genes and are thought to be regulated by their proximal flanking sequences (Delidakis *et al.* 2014). We cloned the 7 kb genomic fragment encompassing these two genes and tagged them with EGFP independently in two otherwise identical genomic constructs: *GFPm7-m8* (EGFP fused to the open reading frame (ORF) of *E*(*spl*)*m7*, **Figure 1 A**) and *m7-GFPm8* (EGFP fused to the *E*(*spl*)*m8* ORF, **Figure 1 B**) (see Supplementary Materials for further features of these transgenes). Consistent with the known *in situ* hybridization expression patterns (de Celis et al., 1996), *GFP-E*(*spl*)*m7* and *GFP-E*(*spl*)*m8* displayed the same pattern in wing imaginal disks from third instar larvae: (*1*) in the region of wing margin (WM), and (*2*) in the adult muscle precursors (AMPs, or adepithelial cells) of the thorax, among other cells (**Figure 1 A, B**). Likewise, both constructs expressed GFP similarly in eye-antennal imaginal disks, whereas their central nervous system (CNS) patterns were different, especially apparent in the Ventral Nerve Cord (VNC) where *GFPm7* was expressed strongly in the midline, while *GFPm8* was mainly expressed in the neuroblasts.

**Figure 1.**
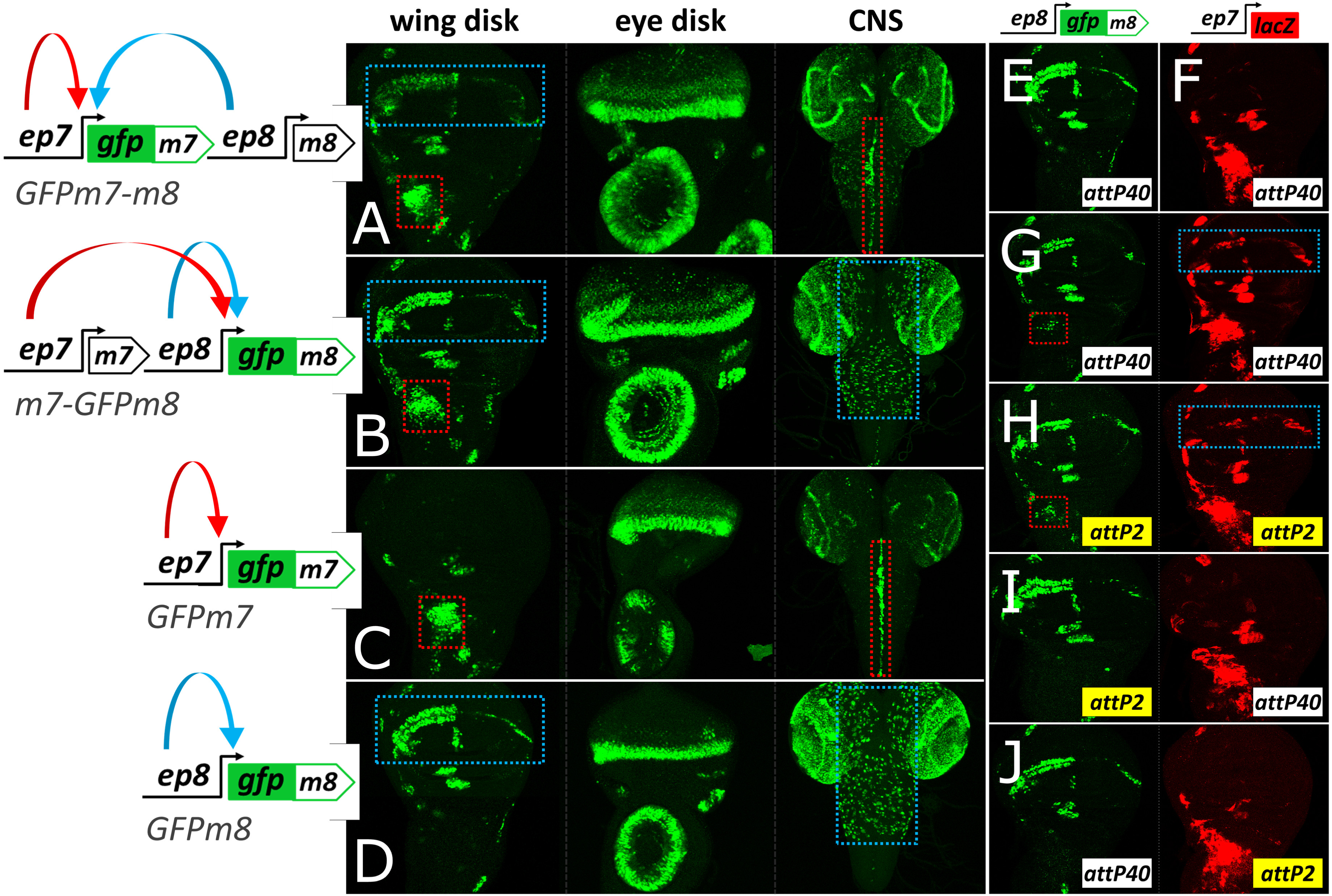
*ep7* and *ep8* interact in *cis* and in *trans*. (**A** and **C**) *GFPm7* and (**B** and **D**) *GFPm8* expression in wing disk, eye-antennal disk and CNS (in left, middle and right columns, respectively) from the EGFP-tagged constructs shown in the diagrams on the left panel; red dotted rectangles highlight e*p7*-specific expression in the adult muscle precursors (AMPs) of the notum and in the midline of the ventral nerve cord (VNC); blue dotted rectangles indicate *ep8*-specific expression in the wing margin (WM) and in neuroblasts of the central brain and VNC. Blue and red arrows in the constructs’ panel depict, respectively, e*7* and *e8* activities which are shared between *p7* (**A**) and *p8* (**B**) in the wing disk. Note lack of WM-specific expression by *ep7* (**C**) and lack of AMP-specific expression by *ep8* (**D**). Also note that both enhancers drive expression in some common areas, e.g. the eye morphogenetic furrow. (**E** and **F**) *cis*-expression patterns in wing disks dissected from hemizygous animals carrying *ep8-GFPm8* (**E**) or e*p7-lacZ* (**F**) transgenes at the *attP40* locus. (**G-J**) *ep8-GFPm8* and *ep*-*lacZ* crossed in the same animal. When present as heterozygotes at the same locus (**G** at *attP40*, **H** at *attP2*) both transgenes expand their pattern to locations dictated by their homologous transgene (marked with dotted rectangles). No such transvection is observed when the *ep8-GFPm8* and *ep7-lacZ* transgenes are placed in the same animal, but at non-homologous loci (**I** and **J**).

We went on to characterize the patterns produced from individual enhancers located immediately upstream of *E*(*spl*)*m7* and *E*(*spl*)*m8* by generating shorter genomic constructs, *GFPm7* and *GFPm8* (**Figure 1 C** and **D**, respectively). *GFPm7* contains *ep7* – the 2.1 kb sequence upstream of *E*(*spl*)*m7* containing its putative enhancer, *e7*, and promoter, *p7. GFPm8* contains *ep8* – the 1.3 kb 5’ flanking *E*(*spl*)*m8* sequence, containing its putative enhancer, *e8*, and promoter, *p8*. *GFPm7* and *GFPm8* recapitulated the expression patterns seen for these genes in the longer *m7-m8* transgenes, with two notable exceptions in the wing disk: *GFPm7* lacked the wing margin (WM) (**Figure 1 C**) and *GFPm8* lacked the muscle precursors (AMPs) (**Figure 1 D**). Additional less conspicuous differences in the bristle proneural clusters and the antennal primordium were noted, but we did not consider these any further. We conclude that these *E*(*spl*) genes contain two upstream enhancers, *e7* and *e8*, which drive distinct expression patterns in the CNS and in the wing disk and similar patterns in the eye disk. In the context of a large genomic fragment encompassing both genes, *e7* and *e8* are shared between promoters of both genes in the WM and AMPs (both genes expressed), but they act exclusively on their downstream gene in the VNC midline (only *m7*) and the neuroblasts (only *m8*).

We also created an *ep7-lacZ* construct to permit simultaneous detection of *ep7*- driven expression of β-galactosidase with *ep8*-driven GFP from *GFPm8* genomic constructs. Both constructs were inserted into the same chromosomal locus via the *ΦC31* integration method (Markstein *et al.* 2008; Pfeiffer *et al.* 2010). *ep7-lacZ* faithfully recapitulated the expression pattern of *GFPm7*. When we made heterozygous animals containing both transgenes (*ep7-lacZ* and *ep8-GFPm8*), we were surprised to find that *ep7-lacZ* displayed novel expression in the wing margin (characteristic of e*8*), while *ep8-GFP* was expressed in the AMPs (characteristic of *e7*) (**Figure 1 G**, compare to **Figure 1 E** and **F**). This effect was observed when both transgenes were inserted into the same attP landing site, either *attP40* (chromosome 2) or *attP2* (chromosome 3) (**Figure 1 G** and **H**). No such inter-transgene interaction was observed when one transgene was inserted in *attP40* and the other in *attP2* (**Figure 1 I** and **J**), suggesting that homologue pairing is required for this interaction; in other words, we are observing a transvection phenomenon.

### Transvection is mediated by homotypic interactions between GIs or Wari insulators

Having encountered robust bi-directional gene activation in *trans* (*ep8* activated by *e7* in the AMPs and *ep7* activated by *e8* at the WM), we decided to dissect the sequences that mediate this phenomenon. Firstly, we asked whether the observed transvection is a general phenomenon requiring only an enhancer paired to a promoter, or whether it depends on the inserted transgene sequences. The constructs in **Figure 1** were based on the *pPelican* vector (Barolo *et al.* 2000), which contains a *pUC8* backbone, the mini-*white* gene, two *gypsy* insulator (GI) sequences flanking a multiple cloning site (MCS), as well as a *ΦC31* attB site inserted by us. In order to systematically address the role of vector sequences we re-cloned the e*p7-lacZ* reporter construct into a minimal *pBluescript* backbone to which we added a *ΦC31* attB site to enable fly transgenesis and either the mini-*white* gene or a *3xP3-dsRed* marker which expresses DsRed in the adult eye in response to an artificial Pax6 (Toy/Eyeless) responsive enhancer (Berghammer *et al.* 1999; Horn *et al.* 2000). All constructs for this analysis were introduced into the *attP40* locus.

Both *pBluescript*-based *ep7-lacZ* constructs (mini-*white* and *3xP3-dsRed*) expressed LacZ in the same pattern as the original *pPelican*-based reporter. However, when they were tested in *trans* to the original *ep8-GFPm8*, no transvection was observed, namely we detected neither LacZ in the WM nor GFP in the AMPs (**Figure 2 A-C**). We concluded that transvection at *attP40* does not happen whenever an enhancer-promoter pair is placed in *trans*, but needs additional sequences from the vectors. Upon flanking the *ep7-lacZ-3xP3-dsRed* reporter with two GIs in the same orientation as in *pPelican*, bidirectional transvection (*e8*→*p7*, blue dotted rectangle and *e7*→*p8*, red dotted rectangle in **Figure 2 D**) was restored implicating the paired GIs as mediators of the effect. Because the GI is known to be bound by the Su(Hw) zinc-finger protein, which mediates its insulator activity (Parkhurst *et al.* 1988; Spana *et al.* 1988; Spana and Corces 1990; Holdridge and Dorsett 1991; Geyer and Corces 1992; Gerasimova *et al.* 1995; Ramos *et al.* 2006), we tested our transgene pairs in a *su*(*Hw*) mutant background. Indeed, *e7*→*p8* (AMP) transvection was lost in this background, but, surprisingly, *e8*→*p7* (WM) transvection was now apparent in all transgene combinations that had a mini-*white* marker in both homologues (**Figure 2 A’, B’** and **E’**), even the one that had not displayed this effect in wild type (wt) genetic background (**Figure 2 B** vs **B’**). We then modified the *pPelican*-based *ep8-GFPm8* construct by removing its GIs. This construct was capable of supporting *e8*→*p7* transvection with the mini-*white* but not the *3xP3-dsRed* version of the *pBluescript*-*ep7-lacZ* reporter in both wt and *su*(*Hw*) genetic backgrounds (**Figure 2 E**/**E’** - **F**/**F’**). The simplest conclusion from these results is that *e8*→*p7* transvection in the WM occurs when mini-*white* is present on both homologues, but this effect is annulled when GIs are placed nearby. Only upon removal of GIs from both homologues, or their inactivation by *su*(*Hw*) loss, is this effect observed. On the other hand, the presence of GIs in both homologues can sustain transvection in both directions, not only *e8*→*p7* but also *e7*→*p8*.

**Figure 2.**
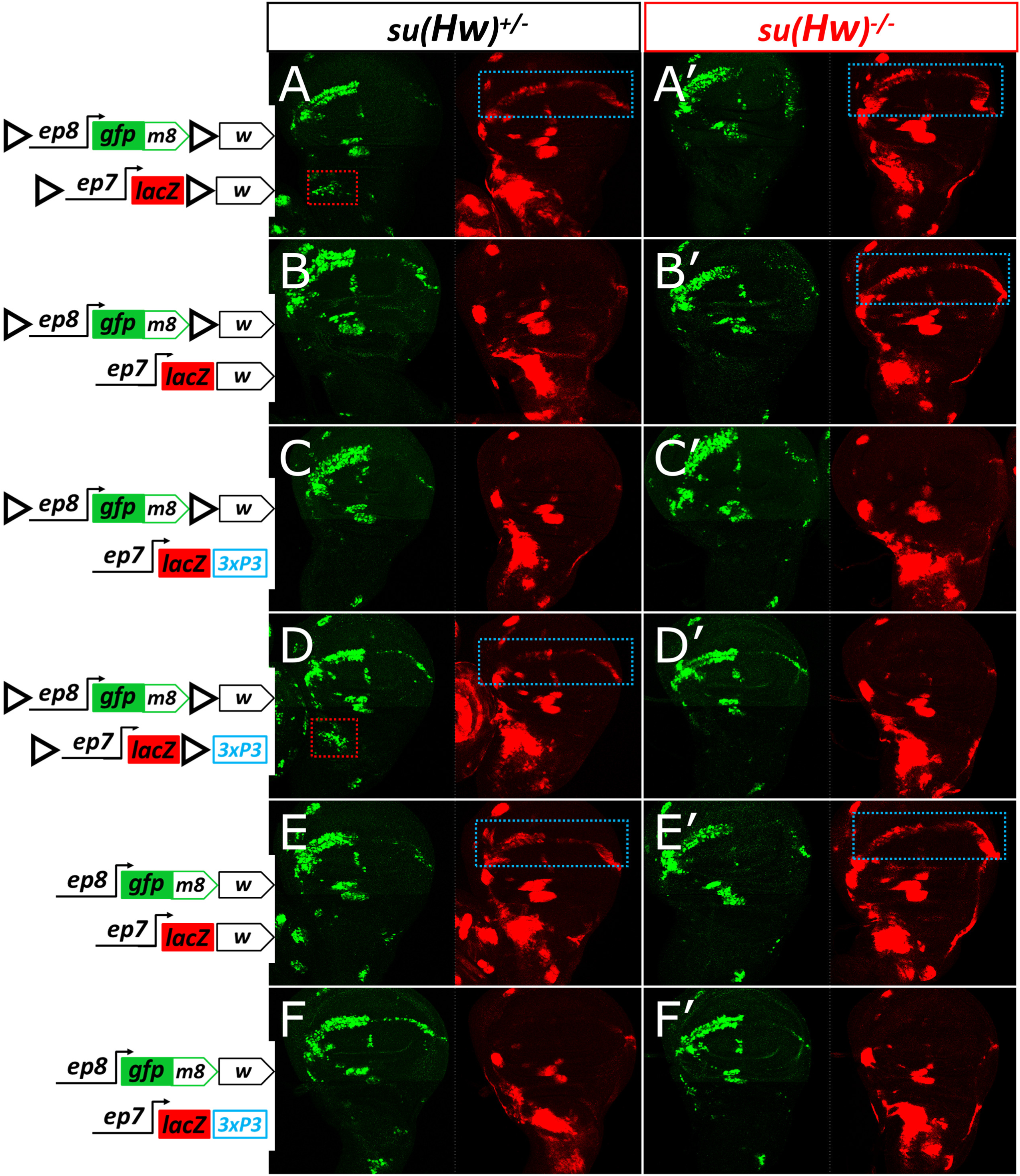
GIs and mini-*white* mediate transvection. Wing imaginal disks from animals heterozygous for variations of the *ep8-GFP-m8* and *ep7-lacZ* transgenes. The two GIs in forward orientation, flanking the two reporter, are indicated by black triangles; marker genes are located downstream of the second (3’) GI: either the mini-*white* gene (‘w’) or the *3xP3-dsRed* cassette (‘3xP3’). An image of the same disk from each genotype is split into two channels: *p8*-driven GFP-m8 in green and *p7*-driven LacZ in red. Dotted rectangles (red and blue) indicate cells (AMPs and WM, respectively) exhibiting transvection. Each combination of transgenes (in rows) is tested in the presence of Su(Hw) (*su*(*Hw*)^*+/-*^, first column, **A**-**F**) and absence of Su(Hw) (*su*(*Hw*)^*-/-*^, second column, **A’**-**F’**). All transgenes are inserted in *attP40* locus.

We mapped the transvection-mediating element of the *white* locus within its 3’ part: out of 3 subfragments derived from mini-*white* in **Figure S1 A-C**, only its 3’most 0.9 kb recapitulated *white-white*-mediated transvection (**Figure S1 C**). It was previously reported that this sequence contains the 3’-UTR as well as an insulator element, dubbed Wari (hereafter referred to WI) (Chetverina *et al.* 2008). Therefore either of two different insulators, GI and WI, can promote transvection when placed in a paired configuration (in both homologues) near an enhancer-promoter pair. WI has been shown to have *su*(*Hw*)-independent insulator activity, but also to interact with GI in *cis* (Chetverina *et al.* 2008). When we confronted a WI-containing *ep7-lacZ* with a WI-containing (mini-*white*) *ep8-GFPm8* flanked by GIs (**Figure S1 D**), transvection was abolished, consistent with what we had observed earlier with the entire mini-*white* (**Figure 2B**). We hypothesized that this inhibition could result from presumptive insulation imposed by the GI located between *e8* and WI in *ep8-GFPm8*, thereby restricting access of WI to *e8*. This was not the case, as the inhibition of WI-mediated transvection was sustained even when we deleted the 3’ GI, leaving only the 5’ GI intact (**Figure S1 E**). Thus, a heterotypic GI-WI interaction in *cis* can disable the homotypic WI-WI-mediated transvection but not the GI-GI-mediated transvection. However, we should emphasize that this inhibition was context-dependent and quite an opposite action of GI, namely enhancement of WI-WI-mediated transvection, was also possible in a different context (described later in this manuscript – see **Figure 8**).

### GI-mediated transvection is promiscuous, whereas WI-mediated transvection is highly selective

Because *ep7* and *ep8* interact in *cis* in their native context (**Figure 1A, B**), it raises the possibility that the transvection we observed is tied to the specificity of these two regulatory modules for each other. Thus, we sought to determine whether GIs and WIs could mediate transvection of *ep7* and *ep8* to an unrelated, heterologous promoter. To address this question, we generated an enhancerless construct containing a basal promoter commonly used in assaying enhancer activity, derived from the *Drosophila hsp70* gene (hereafter referred to as *pH*). As expected, a transgene carrying this minimal *pH* promoter fused to GFP, flanked by GIs and marked with mini-*white* displayed, on its own, no expression in wing disks or CNS (**Figure 3 A**). When crossed to an *ep7-lacZ* construct, congruent in two flanking GIs and mini-*white*, GFP was detected with the *e7*-specific pattern (**Figure 3 B** compare GFP (*trans*) to LacZ (*cis*)). This activation of *pH* in *trans* relies on homotypic interaction between GIs, as disparity between transgenesis markers (but retention of congruent GIs) did not affect transvection of *e7* (**Figure 3 C**), while removal of GIs from one of the transgenes (with congruent mini-*white* markers) abolished the effect (**Figure 3 D**). Unlike the *ep8*→*ep7* transvection, removing GIs entirely, but keeping congruent mini-*white* markers did not support transvection (**Figure 3 H**). Therefore, paired WIs cannot support transvection between *e7* and *pH*.

**Figure 3.**
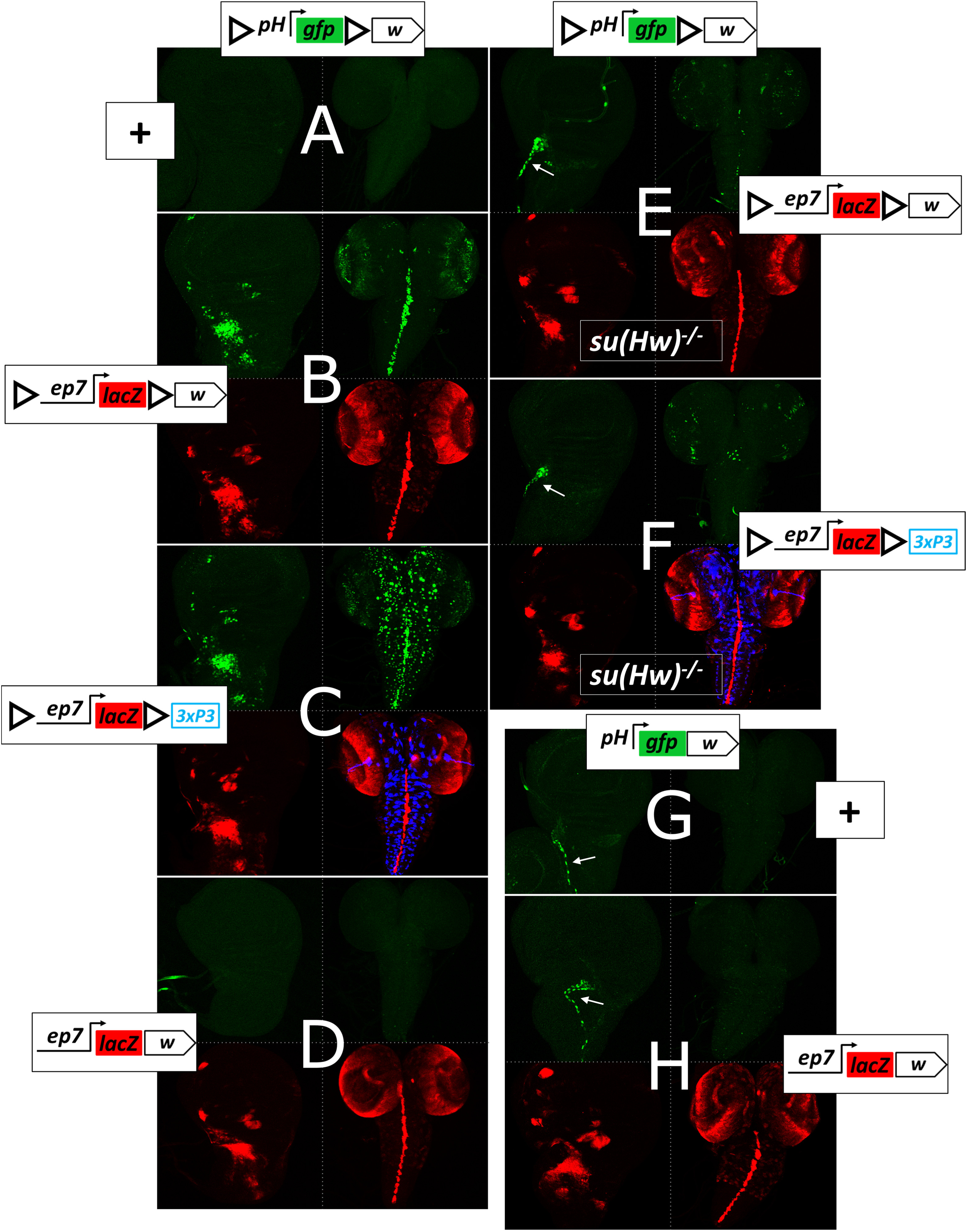
GIs, but not WIs, mediate transvection of *e7* and *3xP3* enhancers to a heterologous, enhancerless *hsp70* promoter (*pH*). Wing disks and CNSs from heterozygotes of *pH-gfp* with GIs (**A**-**F**) or without GIs (**G** and **H**). The second transgene in the homologous locus (*attP40*) is depicted on the side of each panel, the ‘**+**’ sign denotes absence of a second transgene. The *ph-gfp* produces no expression as a hemizygote (**A**). (**B**-**F** and **H**) Projections of *trans* (GFP, green) and *cis* (LacZ, red and DsRed blue, when present) expression from the same wing disk and CNS of each sample containing an *ep7-lacZ* reporter are shown. Note that the observed GFP expression in wing disks of GI-less *pH-gfp* reporters (white arrow) originates from associated tracheal branches in the vicinity of AMPs, and is an effect of trapping a trachea-specific enhancer by the *pH* promoter in *cis* in the absence of the insulating effect of the GIs (**G** and **H**) or their ‘inactivation’ in the *su*(*Hw*)^*-/-*^ background (**E** and **F**). Only combinations of transgenes with GIs in both homologs support transvection (**B** and **C**), while congruency in *white* in the absence of GIs does not (**H**). Consequently, depletion of Su(Hw) protein abolishes transvection observed from **B** and **C** combinations (**E** and **F**, respectively). Note a dotted pattern in the CNS in **C**, manifested by the *3xP3* activity in *cis* (DsRed, in blue) and in *trans* (GFP, in green), which comes from a glial cell population. The artificial 3xP3 enhancer does not drive expression in wing disks.

Interestingly, in this set of experiments, GIs mediated *trans*-activation of the *pH* promoter not only by the *e7* enhancer located in between the two GIs, but also by an enhancer exterior to the two GIs: the *3xP3*, which displays strong expression in a subset of glia in the CNS (**Figure 3C**). Depletion of Su(Hw) suppressed transvection of both enhancers, *e7* (**Figures 3 E, F**) and *3xP3* (CNS in **Figure 3 F**), and allowed *pH* to trap a tracheal enhancer (in *cis*) in the vicinity of *attP40* (white arrows in **Figures 3 E, F**). The fact that two unrelated enhancers, *e7* and *3xP3*, can transvect to a heterologous promoter, *pH*, suggested that GI-mediated transvection is unselective for enhancer-promoter pairs. This prompted us to use the GIs-containing transgene system to screen for putative enhancer elements across the 50 kb long *E*(*spl*) locus. A collection of 18 fragments of this locus inserted in a transgene between two GIs activated specific patterns of expression in *trans* from the GI-flanked *pH-gfp* transgene (**Figure S2**) – a full 15 out of these 18 fragments displayed robust enhancer activity in third instar larval disk/CNS tissue. Additionally, using this system, we were able to recapitulate in *trans* the *cis*-pattern of another enhancer unrelated to the *E*(*spl*) locus, the *vestigial* quadrant enhancer (last column, **Figure S2**).

Unlike GI-, WI-mediated transvection was specific for the *ep8*/*ep7* combination and mediated *trans*-activation of *p7* by *e8* unidirectionally. To test the possibility that WI-mediated transvection is specific for the *E*(*spl*)*m7* promoter (*p7*), we tested various minimal GFP reporters driven by different basal promoters, *pH, p7* and *E*(*spl*)*m8* promoter (*p8*). We confronted these enhancerless non-expressing reporters with an *ep8-lacZ* transgene. We made sure that all combinations were congruent for both GIs and mini-*white*, which enabled us to simultaneously test GI-mediated transvection in a wt background and WI-mediated transvection in the absence of Su(Hw) (**Figure S3**). All three basal reporters responded to *ep8* enhancer in a *su*(*Hw*)^*+/-*^ background (GI-dependent transvection); the *p7* was by far the strongest responder, with *pH* following and *p8* showing a very weak activation (**Figure S3 A-C**). However, in the *su*(*Hw*)^*-/-*^ background (WI-dependent transvection), the *e8* enhancer did not transvect to any of the three promoters (**Figure S3 M**-**O**, compare to **Figure S3 G**-**I**). Therefore, the unidirectional *ep8*→*ep7* transvection supported by WIs was not due to a selectivity of WI for *p7*. When the three basal promoters were fused to the *e7* enhancer (**Figure S3 D-F**), these reporters started expressing (as hemizygotes) in the AMPs and some proneural clusters, but not in the WM, consistent with the activity of the *e7* enhancer (**Figure S3 D-F**). Once confronted in *trans* with *ep8-lacZ*, these *e7*-bearing reporters were able to robustly express GFP in the WM (**Figure S3 J-L**) and this expression was retained in the *su*(*Hw*) mutant genetic background (**Figures S3 P-R**). Thus, the transvection mediated by the interaction between WIs was not promoter context specific, but rather enhancer context specific, with the responding gene requiring the presence of the *e7* enhancer in *cis* in order to sustain WI-mediated transvection. In conclusion, whereas *trans*-paired GIs mediated transvection between any enhancer-promoter pair tested, *trans*-paired WIs were more selective and mediated only *e8*→*e7* transvection, an effect that was dominantly suppressed by the presence of GI elements. This *e8→e7* transvection may reflect some intrinsic affinity of these two enhancers for each other, but it still requires the presence of GIs or WIs in order to materialize.

### Transvection is weaker than cis enhancer-promoter activity and is suppressed by promoter cis-preference

To gauge the relative strength of transvection compared to *cis* enhancer-promoter (e-p) interaction we generated e-p pairs driving GFP expression in *cis* and compared them to the same e-p pairs driven in *trans*. All constructs designed for this purpose were based on the backbone of *pPelican*/*pStinger* vectors (Barolo *et al.* 2000) which contain two GIs to enable GI-mediated transvection, and subsequently inserted into the *attP40* locus. These flanking GIs also provided efficient insulation: all enhancerless promoter-reporter constructs had undetectable levels of expression in all three larval tissues tested (wing disk, eye disk and CNS; data not shown). In all cases we observed lower GFP levels from transvection than from the *cis*-combination (**Figure 4**). We tested three promoters, *p7, p8* and *pH*. Regardless of the enhancer assayed (*e7* or *e8*) the strongest expression levels, both in *cis* and in *trans*, were produced by *p7*, whereas *p8* was the weakest out of the three promoters. This suggests that a promoter’s strength for driving transcription is its intrinsic property and does not depend on the enhancer activating it, at least for the two enhancers tested.

**Figure 4.**
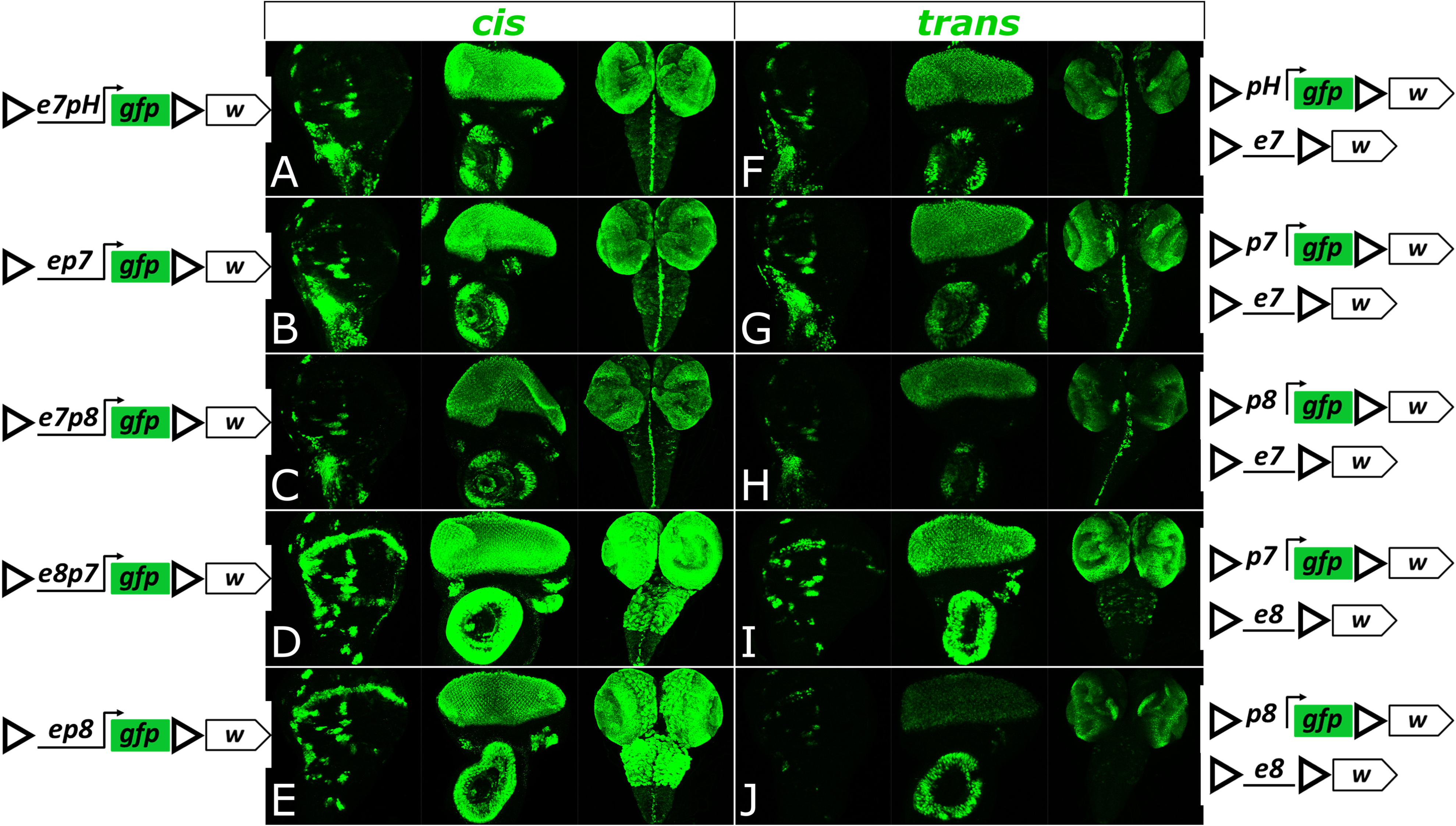
Transcriptional activity of different enhancer-promoter (*e-p*) pairs interacting in *cis* or in *trans* in the presence of GIs. Each genotype is examined for GFP expression in the third instar wing disk, eye disk and CNS. (**A**-**E**) Samples from hemizygotes with a transgene containing a given *e-p* pair linked in *cis* show higher levels of GFP expression than samples from heterozygotes containing same *e* and *p*, in a *trans* configuration (**F**-**J**). All transgenes are inserted in *attP40*.

We also studied the embryonic *cis* versus *trans* expression of *e7* using the *pH* promoter constructs. In *cis, e7* displayed very dynamic embryonic expression, starting at the mesectoderm in stage 7, then in a neuroectodermal cluster pattern up to stage 10 and later in the VNC midline and the epidermis in a complex segmentally repeated pattern. Interestingly, the earlier patterns could not be transvected or were transvected only in a random subset of the cells where the enhancer was active. From stage 11 onward, this sporadic transvection gave way to a more complete one, where the *e7 trans* pattern recapitulated the *cis* pattern (**Figure S4**). This correlates nicely with what is known about somatic homologue pairing: whereas in early embryonic stages paternal and maternal homologues start out unpaired, they gradually increase their pairing and reach maximum levels by about stage 11 (Hiraoka *et al.* 1993; Fung *et al.* 1998; Gemkow *et al.* 1998). Therefore, this observation, corroborates the need for homologue pairing in order for transvection to take place.

To obtain a more quantitative measure of the *cis* versus *trans* activity of an enhancer, we used a luciferase (luc) reporter (instead of GFP) and measured its activity in extracts of larval disk-brain complexes. The GI-flanked *e7-pH-luc* reporter showed 5 times higher activity in *cis* than the *e7* driving an enhancerless *pH-luc* reporter in *trans* (both GI-flanked, **Figure 5 A**). This *trans* activity was still much higher (~26x) than the basal levels of the *pH-luc* reporter. Interestingly, in this assay, *pH-luc* basal levels were low, yet detectable, even though the GFP counterpart had undetectable levels of GFP in the same tissues – this probably reflects the higher sensitivity of the *luc* versus the GFP reporter (Arnone *et al.* 2004). Upon removal of the GIs, *luc* reporter activity dropped to almost undetectable levels, which is consistent with previous observations that GIs can stimulate basal transcription from some promoters (Wei and Brennan 2001; Golovnin *et al.* 2005; Markstein *et al.* 2008; Soshnev *et al.* 2008).

**Figure 5.**
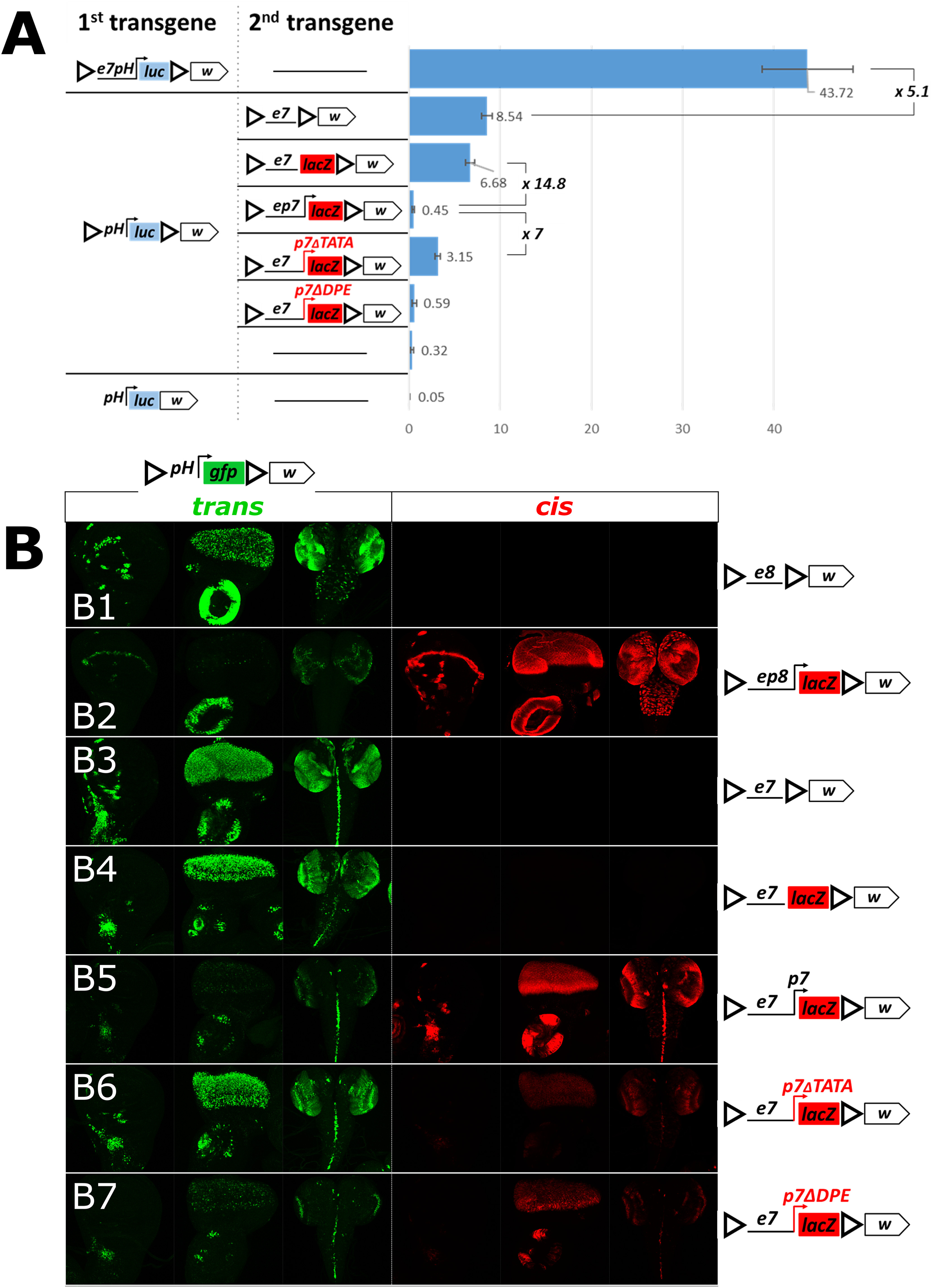
Regulation of GI-mediated transvection by *cis*-preference. (**A**) The chart shows levels of basal and *e7*-induced (in *cis* and in *trans*) *pH*-driven luciferase activity. Levels of luciferase activity were measured from third instar larval disk-brain complexes. Luciferase values normalized to total protein are shown as arbitrary units (a.u.). The mean and standard deviation of 5 replicates is shown. The activity of luciferase transgenes (1st column in the construct panel) is assayed on their own (as hemizygotes; horizontal line in the second column) or in combination with a second transgene in *trans*. (**B1-B7**) Transgenes containing *e8* or *e7*, with or without a *cis*-linked promoter (depicted on the right), were placed in *trans* to *pH-gfp*. For each genotype (each row) the third instar wing disk, eye disk and CNS were examined for (*1*) GFP expression, reflecting *trans* activity of the enhancer-containing transgenes on *pH-gfp* (green) and (*2*) β-galactosidase expression, reflecting *cis* activity of the enhancer-linked promoter, when *lacZ* is present (red). All transgenes contain GIs and mini-*white* and are inserted in *attP40*.

In the above experiments we noticed that confronting an enhancerless reporter in *trans* to a solitary enhancer gave more robust expression compared to all our previous experiments, where the transvecting enhancer was linked in *cis* to a promoter (**Figure 5**, compare **B1** to **B2** for *e8*, and, **B3** and **B4** to **B5** for *e7*). This came as no surprise, since numerous earlier studies on transvection have indicated that an enhancer’s action in *trans* is suppressed by the presence of a promoter in *cis* (Geyer *et al.* 1990; Martínez-Laborda *et al.* 1992; Hendrickson and Sakonju 1995; Casares *et al.* 1997; Sipos *et al.* 1998; Morris *et al.* 1999a; b, 2004; Bateman *et al.* 2012a; Kravchuk *et al.* 2016). Different promoters of varying core element composition have been reported to display *cis*-preference (i.e. to attenuate transvection). As a general rule, mutations compromising transcriptional strength of a *cis*-promoter usually release the enhancer towards *trans* action (Morris *et al.* 1999b, 2004; Lee and Wu 2006). We made two mutations on the *p7* promoter in an attempt to compromise its strength without completely inactivating it. *p7* is a multi-element promoter, containing a TATA box, an initiator (Inr) and a downstream promoter element (DPE) (Klämbt *et al.* 1989; Kutach and Kadonaga 2000). We introduced two deletions into the *ep7-lacZ* construct aiming to disrupt each of these activities; one, *ep7-ΔTATA-lacZ*, removed the TATA box [deletion of −41 to −22 bp relative to the transcription start site (TSS)] and another, *ep7-ΔDPE* -*lacZ*, removed the Inr and DPE elements (deletion of −16 to +67 bp relative to TSS). Both of these promoter mutations retained weak yet detectable transcriptional activity (**Figure 5, B5**-**B7**, *cis* column). Even though the reduction in *cis* promoter activity was comparable between ΔTATA and ΔDPE, the two had dramatically different effects on transvection of the linked *e7* enhancer (**Figure 5, B5**-**B7**, *trans* column). *p7-ΔTATA* partially relieved *e7* from *cis*-preference inhibition, leading to much higher *trans*-activation of *pH-gfp* than that observed with *p7-ΔDPE.* These effects of the mutant *cis*-*p7* promoters on *e7* transvection were independent of the identity of the *trans* promoter, as *p7* and *p8* – based enhancerless reporters responded with a similar trend (**Figure S5**). When the various *ep7-lacZ* versions were confronted with the *pH-luc* reporter, we confirmed that the DPE deletion was comparable to the unmutated promoter in strongly suppressing transvection (11-15x weaker than a promoterless *e7-lacZ*), whereas the TATA deletion released *e7* from *cis*-preference giving 6-7x stronger *trans* reporter expression (compared to the unmutated e*p7* or the e*p7ΔDPE*) (**Figure 5 A**). Interestingly, in this assay the intact e*p7* (and e*p7-ΔDPE*) produced very low *trans*- activation of the *pH-luc* reporter, only 1.4-2x higher than its basal levels attained in the absence of a transvecting enhancer. We speculate that this reflects the ability of the transvecting *e7* enhancer to activate *pH-luc* in a number of cells (as visualized by the *pH-GFP* reporter) but at the same time to *repress* the basal *pH-luc* activity in the remaining cells. Unfortunately, we can only measure the resultant luciferase activity in the whole brain-disk extract with no cell-to-cell resolution, so we cannot test this scenario. Regardless, these luciferase constructs enabled us to obtain a quantitative measure of transvection strength, which ranged from 5x to almost 100x lower than the *cis* output of the same promoter-enhancer pair, depending on the presence of a *cis*-linked promoter and in particular the integrity of its TATA box.

### Relative position, number and orientation of GIs determine transvection effects

In all the previous experiments, all our GI-based transgenes contained two GIs each in forward orientation (GIs^FOR^) as in the *pPelican* and *pStinger* series vectors: the ‘5’ GI^FOR^’ at the 5’ end of each construct (following only the *φC31* attB integration site), and the ‘3’ GI^FOR^’, 3’ to the *lacZ* or *gfp* reporters (preceding the *3xP3* or mini-*white* marker genes, **Figure S6**). The “forward” GI orientation is the same as the one found in the original *gypsy* transposon, where GI is located shortly downstream of the 5’ LTR (ref). The two transgenes (*ep7-lacZ* and *pH-gfp*) in this starting configuration of GIs (hereafter referred to as the ‘dual-GIs^FOR^’ configuration), when presented to each other in *trans*, result in expression of GFP in two distinct patterns: *e7*-specific in all tested tissues (wing disk, eye disk and CNS) and *3xP3*-specific in the CNS (**Figure 3** and **Figure S6**). Thus, the *pH* located in between the two GIs^FOR^ receives input from two enhancers in *trans*: *e7* – located downstream of the 5’ GI^FOR^, and *3xP3* located downstream of the 3’ GI^FOR^. We have introduced a series of modifications in the configuration of the GIs within these two constructs in order to understand how the relative number, position and orientation of the GIs influence transvection. All resultant constructs for this analysis were introduced into the *attP40* locus.

Deletion of the 3’ GI^FOR^ in the transvection-receiving construct, *pH-gfp*, while preserving the dual-GIs^FOR^ configuration in the ‘sending’ *ep7-lacZ* construct, caused a reduction in transvection of *e7*, with concomitant increase of *3xP3* transvection (compare **Figure 6 A** to **B**). When the 3’ GI^FOR^ was deleted in the ‘sending’ construct, and the ‘receiving’ construct was kept in its initial dual GIs configuration, the transvection of *e7* seemed unaffected, while transvection of *3xP3* was nearly lost (compare **Figure 6 C** to **A**). Finally, deletion of the 3’ GIs^FOR^ in both constructs led to augmented GFP expression with an *e7* pattern and an almost undetectable *3xP3* pattern (compare **Figure 6 D** to **A**). These data demonstrate that a robust *trans*- activation of the *pH* promoter by the *e7* enhancer is mediated via an interaction between the 5’ GIs^FOR^ of the two transgenes. This interaction seems to be weakened by the presence of a 3’ GI^FOR^ in either or both of the interacting transgenes. However, the 3’ GI^FOR^ in the sending construct is required for effective transvection of the *3xP3*, suggesting that the *3xP3* enhancer, like *e7*, needs an adjacent GI in order to robustly act in *trans*.

**Figure 6.**
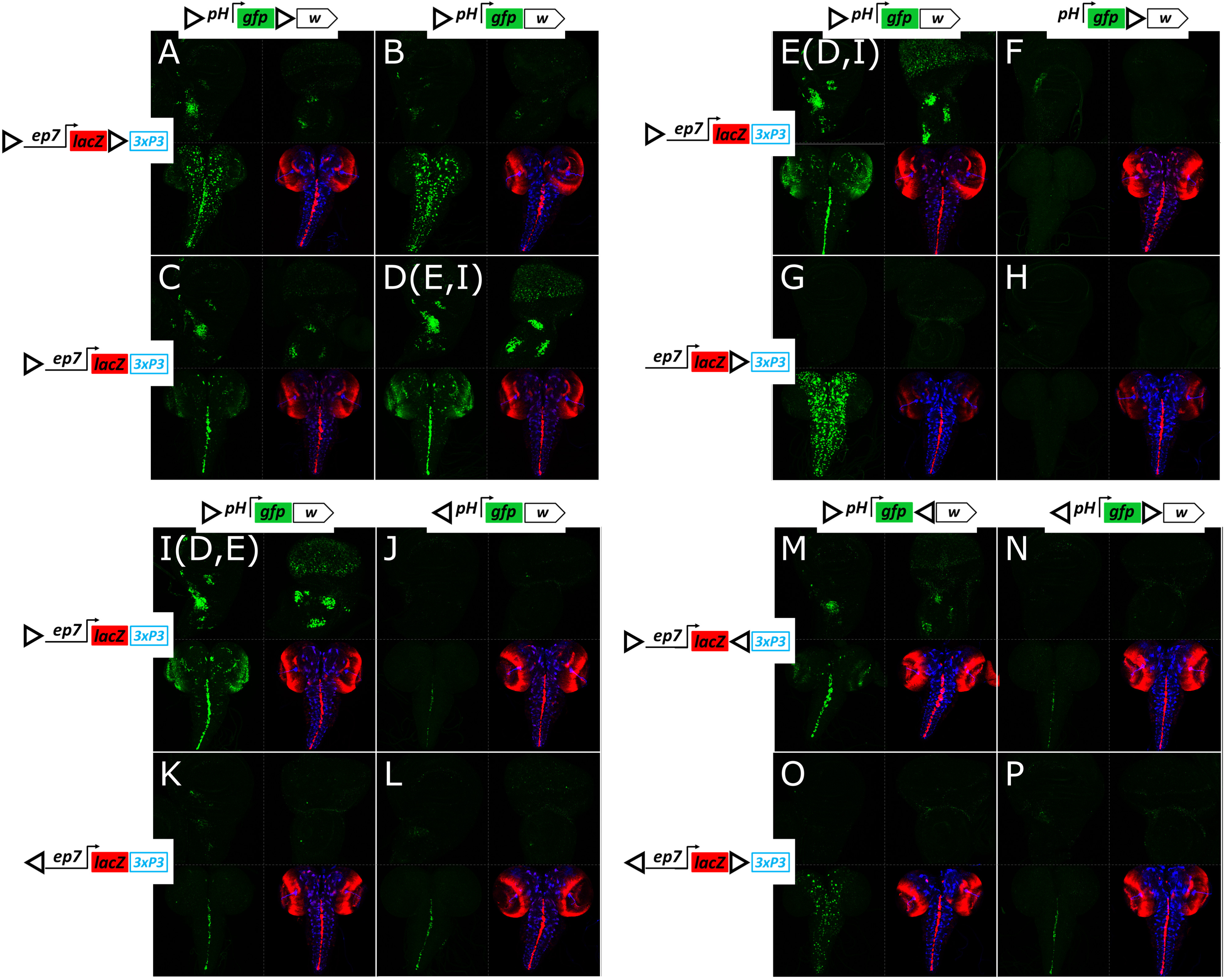
The relative position, number and orientation of GIs determine transvection effects. (**A**-**P**) Confocal z-projection of GFP expression in third instar larval wing disk (top left panel for each genotype), eye disk (top right) and CNS (bottom left). Merged *ep7*-driven LacZ (red) and *-3xP3*-*pH-*driven, DsRed expression (blue) in the same CNS (bottom right); note that the *3xP3* reporter shows no expression in the third instar eye or wing disk; for patterns of *ep7-lacZ* expression in disks, see previous figures. Each genotype contains a *pH-gfp* transgene with mini-*white* in trans to an *ep7-lacZ* transgene with *3xP3*-dsRed and various arrangements of GIs, as indicated. Note, that **D, E** and **I** represent different samples obtained from the same genotype. All transgenes are inserted in *attP40*.

Consistently, when the dual-enhancer construct (***e****p****7****-lacZ-****3xP3****-dsRed*) contained a single GI^FOR^, only its downstream enhancer was transvected to a GI^FOR^–preceded *pH-gfp*: *e7* (with weak sporadic activity of *3xP3*) (**Figure 6 D**,**E**,**I**) or *3xP3* (**Figure 6 G**). Moreover, the presence of the GI^FOR^ upstream of the *pH* promoter was essential for its activation by a *trans*-enhancer, as its deletion abolished transvection altogether, even when another GI^FOR^ was present at the 3’ position (**Figures 6 F** and **H**). It therefore seems that the *trans*-activity of both enhancers (*e7* and *3xP3*) obtained from the dual-GIs^FOR^ *ep7-lacZ* transgene (**Figures 6 A** and **B**) resulted from the interaction between the 5’ GI^FOR^ preceding *pH* with the two GIs^FOR^ of *ep7-lacZ* in *trans*, each upstream of each enhancer.

We next addressed the significance of the orientation of GIs in mediating transvection. Surprisingly, the presence of a reversed GI in the 5’ position (5’ GI^REV^) in either of the two transgenes strongly reduced transvection effects, even when both transgenes contained 5’ GI^REV^ (**Figure 6 J-L** compared to **I**). Therefore, *trans* interaction between enhancers and promoters is favored when both are located on the 3’ side of GI. Weak transvection, on the other hand, can be sustained regardless of 5’ GI orientation. In fact, even incongruent combinations of 5’ GIs (one FOR and the other REV, **Figure 6 J, K**) displayed weak transvection, suggesting that it is not so much the congruence of the *trans*-insulator pair, but rather the absolute orientation of both GIs that is needed for robust transvection.

The fact that the preferred position of the *trans*-interacting enhancer-promoter is on the 3’ side of the two GIs made us consider the possibility that placing the 5’ and 3’ GIs in a convergent orientation (i.e., 5’ GI^FOR^/ 3’ GI^REV^) in both constructs might strengthen *trans*-interaction. However, this was not the case, as such transgenes produced equal levels of transvection to those with 5’ and 3’ GIs in the forward orientation (**Figure 6 M** compared to **A**) and less than the combination where 3’ GIs are absent altogether (**Figure 6 D).** These results suggest that the interaction between 5’ GI^FOR^-preceded enhancer and *trans*-promoter is weakened by the presence of a second (3’) GI in *cis*, irrespective of its orientation. Moreover, transgenes with a divergent configuration of GIs (5’ GI^REV^/ 3’ GI^FOR^) did not improve the weak transvection observed between transgenes with a single 5’ GI^REV^ (compare **Figure 6 P** to **L**), nor were incongruent 5’ GI configurations improved by a 3’ GI (**Figure 6 N, O**) (note, however, that *3xP3* was efficiently transvected to *pH* in **Figure 6 O** as both *3xP3* and *pH* are preceded by GI^FOR^).

Is the ability of a single 5’ GI^FOR^ to support transvection a peculiarity of the attp40 locus or can it happen in more genomic loci? When we tested constructs with a single 5’ GI^FOR^ in four more *attP* loci and we got robust transvection in all cases (**Figure S7**). Importantly, removing the 5’ GIs^FOR^ from the transgenes abolished transvection in all loci, reconfirming the need for paired homotypic insulators in both homologues. As a corollary, we conclude that, at least in the five genomic loci we tested, nearby endogenous genomic insulators were not capable of mediating transvection from the GI-less constructs, suggesting that “insulator trapping” is probably not a common phenomenon in the *Drosophila* genome. On the contrary, enhancer trapping is very common; we were able to detect some non-*e7* dependent patterns of expression of our transgenes in four out of the five loci tested (all except *VK40*; see **Figure S7**).

In summary, transvection needs homotypic insulators in both homologues, but having homotypic insulators is not sufficient. The outcome is also influenced by the *position, orientation* and *number* of these insulators. In the context of the *ep7-lacZ-3xP3* → *pH-gfp* transvection, both GIs have to be 5’ of the *pH* promoter and directly adjacent to the transvecting enhancer, *e7* and/or *3xP3*. The FOR orientation is greatly favoured for both homologues; the REV orientation produces a much weaker effect. Finally adding another GI in one or both transgenes weakens the 5’GI^FOR^ mediated transvection. It should be kept in mind, however, that the transcriptional outcome of a *trans*-interacting insulator pair, is also greatly dependent on the enhancers and promoters located in the vicinity of these GIs: when the same *ep7-lacZ* series of transgenes was tested in *trans* to an *ep8-m8GFP* series (instead of the *pH-GFP*), the *ep7*→*ep8* transvection largely obeyed the above rules, but the *ep8*→*ep7* transvection was detectable even with a single 3’ GI, regardless of orientation (**Figure S8**). Still, the single 5’ GI^FOR^ configuration gave the strongest *trans*-effect even with this transgene combination. The removal of GI from either homologue completely abolished the effect, as expected.

Why was the addition of a 3’ GI^FOR^ detrimental to *ep7*→ *pH* transvection (**Figure 6 A**- **D, M-P**)? Could it potentially engage in homotypic interactions with either the *cis* or the *trans* 5’ GI that might compete with the ability of these 5’ GIs to support transvection? To gain insight on the activity of this 3’ GI^FOR^ we appended a “tester” module to our GI^FOR^-*ep7-lacZ-*GI^FOR^-*3xP3-dsRed* transgene. Two different modules were cloned immediately 3’ of the 3’ GI^FOR^: (*1*) a ‘receiving’ *pH-gfp* module or (*2*) a ‘sending’, *ep8-m8* module (*ep8* driving an untagged *E*(*spl*)*m8* CDS). **Figure 7** presents the results obtained in wing disks, which are consistent with those obtained in the CNS and eye-antennal disk (data not shown). The tester modules did not influence *ep7-lacZ* expression: both transgenes, on their own, expressed LacZ in the AMPs, as expected (**Figure 7 A**/**A’**); an *e8*-specific wing margin LacZ pattern was not detected, consistent with insulation of *ep7-lacZ* from *e8* (**Figure 7 A**); similarly, the transgene containing the insulated enhancerless *pH-gfp* module showed no GFP expression (**Figure 7 A’)**, suggesting that the 3’ GI in this construct insulates *ep7* from *pH*, instead of enabling their interaction. It therefore seems that the two GIs^FOR^ in these tester constructs do not productively interact in *cis* (see also Cai and Shen 2001; Kyrchanova *et al.* 2008a). When tested against each other, we observed a robust activity of the *pH-gfp* module in the WM, indicative of a *trans* interaction between the two 3’ GIs^FOR^ resulting in *ep8-m8*→*pH-gfp* transvection (**Figure 7 B**). Therefore, 3’ GIs prefer to homotypically interact in *trans*. The presence of only few GFP positive AMPs and no apparent WM LacZ expression in this combination (**Figure 7 B**) demonstrate that the “diagonal” interactions between GIs^FOR^ (i.e., 5’ GIs^FOR^ −3’ GI^FOR^) are less favored than the “vertical” ones (5’-5’ and 3’-3’).

**Figure 7.**
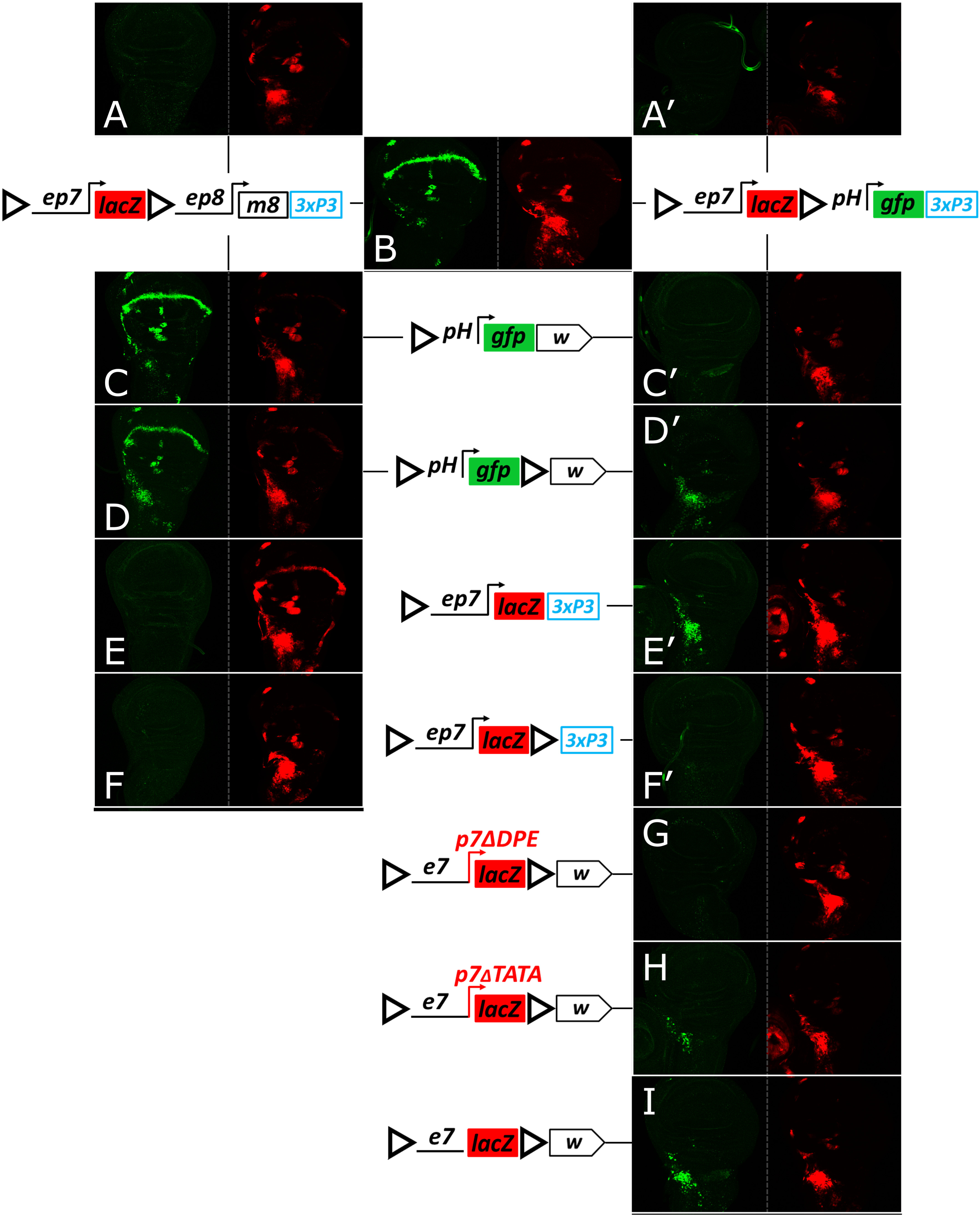
In dual GI transgenes, both 5’ and 3’ GIs participate in trans-interactions. Two tester transgenes, GI^FOR^-*ep7-lacZ-*GI^FOR^*-ep8-m8* (left column, **A**-**F**), and GI^FOR^-*ep7-lacZ-*GI^FOR^-*pH-gfp* (right column, **B, A’**-**F’, G**-**I**) were tested as hemizygotes (**A, A’**), combined *inter se* (**B**) or combined with various other GI-containing transgenes in *attP40* (**C**-**I**). Third instar wing imaginal disks are shown for each genotype in two channels (GFP in green, LacZ in red). Note stronger *e7*→*pH-gfp* transvection in **D** and **D’** compared to **C** and **C’**. Also note that the ability of the 3’ GI^FOR^ of both tester constructs to interact with a 5’ GI^FOR^ of the *ep7-lacZ* reporter (**E, E’**) is severely compromised when a second GI is added to the *ep7-lacZ* reporter (**F, F’**). Interestingly this suppression is relieved by disabling the *p7* promoter in the latter construct (**G**-**I**).

Placing the 5’ GI^FOR^ *pH-gfp* transgene in *trans* to the testers gave strong transvection of *e8* (via the 3’ GI^FOR^; **Figure 7 C**) but only weak or no transvection of *e7* (via the 5’ GI^FOR^; **Figure 7 C, C’**). Interestingly, *e7*→*pH* transvection with both testers was augmented when a 3’ GI^FOR^ was added to the responding *pH-GFP* transgene, as evidenced by broadly expressed GFP in the AMPs (**Figure 7 D, D’**). Therefore, when confronted with dual GIs^FOR^, a single GI^FOR^ preferentially interacts with the *trans* 3’ GI^FOR^, but this shifts to the 5’ GI^FOR^ when a second GI^FOR^ is added resulting in a dual/dual configuration. This conclusion was supported by confronting the tester transgenes with a single vs dual GI^FOR^ configured *ep7-lacZ* reporter. Only the single 5’ GI^FOR^ was able to interact in *trans* with the two tester modules, as evidenced by *ep8-m8*→*ep7-lacZ* and *ep7-lacZ*→*pH-gfp* transvection (**Figure 7 E, E’**), whereas both of these transvection effects were lost in the dual/dual configuration (**Figure 7 F, F’**). Surprisingly, the *ep7-lacZ*→*pH-gfp* transvection was regained when the *p7* promoter was compromised or (better) deleted in the dually GI^FOR^ flanked *ep7-lacZ* transgene (**Figure 7 G**-**I**). We therefore conclude that GIs sample different homotypic interactions in *trans*, both vertical and diagonal. The latter are disfavored, but can still occur and their ability to support transvection is influenced by the enhancer-promoter interactions in their vicinity, e.g. the *ep7-pH* is not sufficient to produce diagonal transvection (**Figure 7 F’**), unless relieved from *cis* promoter preference (**Figure 7 H, I**). As another example, a similar diagonal GI-GI interaction can produce transvection of *e8* (from the *ep8-m8* tester) to *pH-gfp* (**Figure 7 D**), but not to *ep7-lacZ* (**Figure 7 F**). Such alternative *trans*-interactions in dual-GI combinations probably compete with 5’GI - 5’GI interactions and could underlie the suppression of transvection produced by the addition of 3’ GIs in **Figure 6 A-D**.

Finally, we note that the *3xP3*→*pH* transvection in **Figure 6 A** is also mediated by a 5’ – 3’ (diagonal, less favored) GI^FOR^ interaction. We tested the same configuration of enhancers and insulators and changed only the responding promoter on the dual GIs^FOR^ *p-gfp* construct. Only *pH* was able to strongly respond to *3xP3*, whereas two other promoters, *p7* and *p8*, showed no or very weak response (**Figure S9**), consistent with an important role for enhancer-promoter combination to bias the transvection outcome of a less favorable GI-GI homotypic interaction.

### Other insulators also mediate transvection

To determine whether other *Drosophila* insulators also mediate transvection, we generated two series of constructs based on the *pLacZattB* vector (Bischof *et al.* 2007): ‘sender’ constructs containing an insulator cloned in between the *e7* and *e8* enhancers, and ‘responder’ constructs containing an insulator between divergently oriented *pH-lacZ* and *pD-gfp*, a reporter driven by the *Drosophila Synthetic Core Promoter* (DSCP, abbreviated as *pD*) (Pfeiffer *et al.* 2008). Besides GI, we tested two new insulators: (*1*) the Fab8 insulator isolated from the *Abdominal-B* region of the *bithorax* complex; Fab8 insulator activity depends on dCTCF, the ortholog of the vertebrate CTCF protein (Barges *et al.* 2000; Moon *et al.* 2005; Kyrchanova *et al.* 2008b, 2016), and (*2*) 1A2, an insulator located downstream of the *yellow* gene, containing two Su(Hw) binding sites (Golovnin *et al.* 2003; Parnell *et al.* 2003). Fab8 and 1A2 exemplify two major classes of endogenous insulators [defined by binding of dCTCF and Su(Hw), respectively] which are abundantly represented in the *Drosophila* genome (Parnell *et al.* 2006; Adryan *et al.* 2007; Negre *et al.* 2010; Baxley *et al.* 2017). In addition, the resultant constructs contain WI carried in the mini-*white* marker gene. All transgenes were inserted into the *attP40* locus and we present the results from wing disks, which are consistent with the results obtained from the CNS and eye-antennal disks (not shown).

The responder transgene without an insulator between *pH* and *pD* promoters (“blank” responder), on its own, showed trachea-specific activity of both promoters (both LacZ and GFP, **Figure 8 A1**), similar to the expression of the uninsulated *pH-gfp* reporter at the *attP40* locus (**Figure 3 G**). Inserting GI between the two promoters insulated *pD-gfp* from the tracheal enhancer and resulted in the trapping of (an)other enhancer(s) at the *pD* promoter, ubiquitously active in all cells of the disk’s epithelium (weak ubiquitous GFP expression in **Figure 8 A2**). This latter, ubiquitous, activity was abolished by a deletion of the WI from the mini-*white* (**Figure 8 A3**) and was not observed with WI on its own (**Figure 8 A1**) indicating cooperation between GI and WI in mediating the *cis* activity of this enhancer onto *pD*. The Fab8 responder, similarly to the GI responder (**Figure 8 A2**), also trapped the epithelial enhancer via the *pD* promoter (**Figure 8 A4**), while the 1A2 responder did not show any activity from any of the promoters (**Figure 8 A5**).

**Figure 8.**
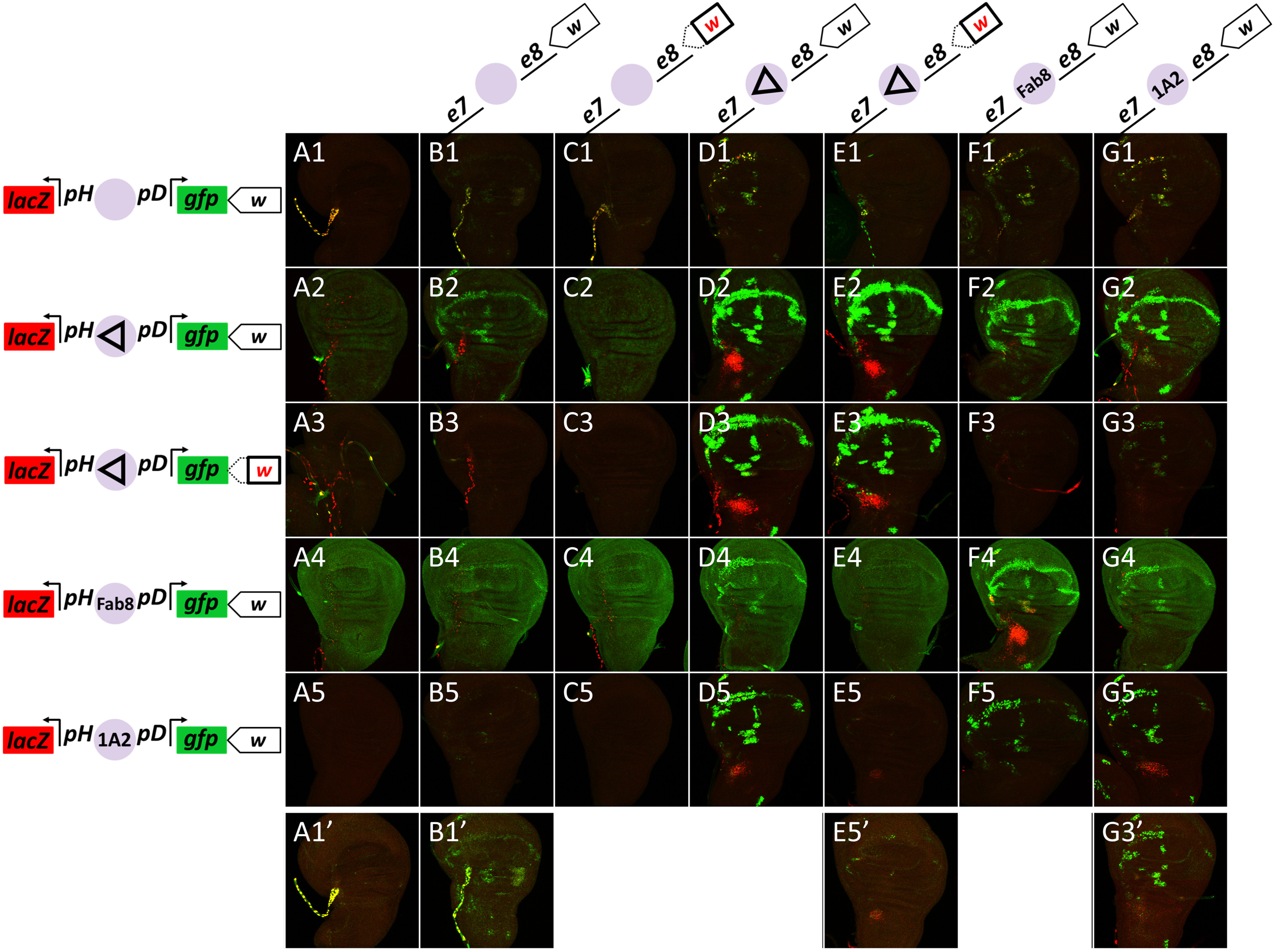
1A2 and Fab8 insulators also mediate transvection. (**A1**-**G5**) Merged GFP and LacZ channels of confocal z-projections from third instar wing disks. (**A1**-**A5**) Wing disks from animals hemizygous for responder transgenes. (**B1**-**G5**) Wing disks from heterozygotes for sender transgenes (as indicated in each column) with responder transgenes (as indicated in each row). All transgenes are inserted in *attP40*. **A1**-**G5** are imaged at the same intensity settings. **A1’**, **B1’**, **E5’** and **G3’** are higher-sensitivity images of the respective panels, to reveal very low levels of transvection. Note that the responder construct in row **3**, as well as the sender constructs in columns **C** and **E**, are deleted for WI.

Heterozygotes between the “blank” sender and “blank” responder transgenes produced extremely faint but visible expression of GFP in the WM, indicative of a WI-mediated *trans* interaction between *e8* and *pD* (**Figure 8 B1, B1’**). Surprisingly, this interaction was augmented in the GI responder (WM in **Figure 8 B2**). This enhancement of WI-WI mediated transvection by GI, was confirmed by deleting one WI, which abolished GFP expression in the WM (**Figure 8 B3**). This is in contrast to our previous result where GI-WI interaction in *cis* had an inhibitory effect on the WI-mediated transvection (**Figure 2**), suggesting that the transcriptional outcome of GI-WI cis interaction is context-dependent. Fab8 also showed a detectable, albeit weaker, enhancement of *e8*→*pD* transvection at the WM, whereas 1A2 had no detectable effect (**Figure 8 B4, B5**). Consistent with the conclusion that this transvection was mediated by homotypic WI/WI, when we deleted WI from the sender transgene, no transvection was observed at the WM in combination with any of the responders (**Figure 8 C1-C5**). WI-mediated transvection was also weakly enhanced by adding a heterologous insulator in the sender homologue (**Figure 8 D1-G1**) and testing against the “blank” responder. Again, only *e8* was transvected, although this time both *pH* and *pD* responded, consistent with the fact that no insulator lies between the two. In this case 1A2 was able to enhance transvection comparably to GI and Fab8 (**Figure 8G1**, compare to **D1** and **F1**). Removal of WI from the GI sender abolished transvection (**Figure 8E1**).

Unlike the weak transvection effects discussed so far with the “blank” senders (**Figure 8**, column B) or responders (**Figure 8**, row 1), where only WI was present, we got very strong transvection of both *e7* and *e8*, when we combined homotypic insulators in sender and responder, i.e., GI/GI, Fab8/Fab8 or 1A2/1A2 (**Figure 8 D2, D3, E2, E3, F4, G5**). In all cases *e7* transactivated the *pH-lacZ* reporter (in the AMPs) and *e8* transactivated *pD-gfp* (in the WM), consistent with orientation-dependent function of all three insulators. GI produced the strongest effect and 1A2 the weakest. When the WI was deleted from either the sender or the responder GI construct, no difference in transvection efficiency was seen (**Figure 8 D2, E2, D3, E3**) thus confirming, in a different context, our earlier conclusion that WIs do not affect GI-mediated transvection.

Moderately strong WM GFP expression (*e8*→*pD* transvection,) was also seen in apparently heterotypic insulator combinations, specifically Fab8 or 1A2 senders with a GI responder (**Figure 8 F2, G2**) and the reciprocal, i.e., a GI sender with Fab8 or 1A2 responders (**Figure 8 D4, D5**). Upon deleting the WI from either the GI sender or responder, however, all of these effects were abolished (**Figure 8 F3, G3, E4, E5**), consistent with being mediated via WI/WI and enhanced by the presence of the heterologous insulators, similar to the effects noted earlier with “blank” sender/ responder constructs. On the other hand, the AMP *lacZ* and WM GFP expression seen in both 1A2/GI combinations (**Figure 8 G2** and **D5**) was maintained, albeit much more weakly, upon WI deletion (**Figure 8 G3**/**G3’** and **E5**/**E5’**), suggesting that this results from a *trans* heterotypic interaction between GI and 1A2, which is not surprising, since both are Su(Hw)-binding insulators.

In conclusion, we provide evidence that insulator landscape can affect enhancer-promoter communication both in cis (enhancer trapping) and in trans. All homotypic insulator combinations tested support transvection. The presence of additional heterotypic insulators in one or both homologues can augment (or in other contexts suppress) this effect. We finally provide evidence that heterotypic insulators can weakly promote transvection if they belong to the same class.

## Discussion

Transvection is the ability of an enhancer to activate transcription from an unlinked promoter located at the same locus of the homologous chromosome. Using a collection of enhancers and promoters driving GFP and LacZ reporters and targeted to specific genomic loci, we were able to study transvection and characterize important parameters influencing its outcome. The salient features of this phenomenon borne out by our results are the following: (1) We confirmed that homologue pairing is a prerequisite for transvection, as already known from classical studies. In *Drosophila*, homologue pairing in non-meiotic cells is widespread, but seems to evolve gradually during the first half of embryogenesis (Hiraoka *et al.* 1993; Fung *et al.* 1998; Gemkow *et al.* 1998) – we showed that transvection unfolds in a similarly gradual manner, being stochastic and erratic during the early stages of embryogenesis, while recapitulating precisely the *cis*-activity of the enhancer at later embryonic and larval stages, once homologue pairing has been completed. (2) Insulators are needed for transvection. At least four different insulators, GI, WI, Fab8 and 1A2, are capable of mediating transvection. We focussed on GI and WI, which are commonly found in transgenesis vectors. (3) GI-mediated transvection is strong, resulting in about 20% of the expression level the same enhancer-promoter combination would give in a *cis* configuration, and can work with all enhancer-promoter combinations tested. In contrast, WI-mediated transvection is weak and is only detectable if the responding promoter is accompanied by another enhancer (**Figure S3**) or another insulator (**Figure 8 B2**). (4) While necessary, the presence of GIs in both homologues is not sufficient for transvection. (4a) The most important parameters in determining the transvection outcome are the number and position of GIs in both homologues – the strongest effects are seen when a single GI is present in each homologue, one immediately upstream of the responding promoter and another adjacent to the interacting enhancer. Additional GIs at other positions of homologously inserted transgenes have a detrimental effect, probably by competing against the 5’GI/5’GI enhancer-promoter enabling interaction. (4b) The presence of a promoter in *cis* to the enhancer reduces its effectiveness in transvection. This *cis*-preference phenomenon, which has been described before (Geyer *et al.* 1990; Martínez-Laborda *et al.* 1992; Hendrickson and Sakonju 1995; Casares *et al.* 1997; Sipos *et al.* 1998; Morris *et al.* 1999a; b, 2004; Bateman *et al.* 2012a; Kravchuk *et al.* 2016), depends on the activity of the *cis*-linked promoter: we have shown that the TATA element seems to play a more important role than the DPE in inhibiting a *cis*-linked enhancer from acting in *trans*. (4c) The GI is highly asymmetric with 12 Su(Hw) binding sites all in the same orientation: the GI orientation with respect to the enhancer-promoter interacting pair is also important, although not crucial for the transvection outcome. The 3’ side of the GI is the optimal for promoting transvection, but the 5’ side is also functional but less effectively. (5) Non-homotypic insulators generally do not promote transvection, with the exception of GI/1A2, both of which bind Su(Hw) and showed a weak interaction. (6) Non-homotypic insulators do however *cis*-influence the transvection produced by homotypic insulators in a context-dependent manner. For example GIs can enable or inhibit WI/WI-mediated transvection. However, WI seems unable to affect GI/GI-mediated transvection.

A role of insulators in transvection has been described, but the mechanism been at best unclear (Fukaya and Levine 2017). Whereas some studies propose that insulators are needed for transvection (Lim *et al.* 2018), others propose that they only have an accessory role (Kravchenko *et al.* 2005; Schoborg *et al.* 2013), or that they affect transvection by promoting homologue pairing (Fujioka *et al.* 2016). And finally, other studies ignore them altogether (Bateman *et al.* 2012a; Mellert and Truman 2012; Blick *et al.* 2016). Instead, the classical proposed role of insulators is to insulate, i.e. to inhibit enhancer-promoter communication in *cis*, although sometimes they can enable such enhancer-promoter communication, e.g. via their so-called “bypass” activity (Cai and Shen 2001; Muravyova *et al.* 2001; Kyrchanova *et al.* 2008a; Fujioka *et al.* 2013). These two apparently contradictory activities have been reconciled by the “looping” model (reviewed in Chetverina *et al.* 2017; Schwartz and Cavalli 2017), which is based on evidence that insulators mediate homotypic, or sometimes heterotypic, interactions (Kyrchanova *et al.* 2008a; Li *et al.* 2011; Vogelmann *et al.* 2014; Bonchuk *et al.* 2015; Fujioka *et al.* 2016). Via these interactions, insulators can form chromatin loops and these loops can either bring enhancers and promoters in proximity (e.g. when both are near the loop’s anchor points) or avert their proximity (e.g. when one is within one loop and the other is outside that loop). These activities occur in *cis* and shape chromosomal architecture in parallel to affecting transcriptional regulation. The same insulator-insulator interactions can occur in *trans* (Kravchenko *et al.* 2005; Fujioka *et al.* 2016; Lim *et al.* 2018) and this could mediate interactions of enhancers on one homologue with promoters on the other (transvection).

One model proposes that insulators promote transvection by mediating homologue pairing in somatic cells. This hypothesis is supported by the fact that different classes of insulators are widely distributed in the *Drosophila* genome (Bartkuhn *et al.* 2009; Bushey *et al.* 2009; Negre *et al.* 2010; Schwartz *et al.* 2012) and their congruent matching could underlie paternal-maternal homologue alignment from end to end. Alternatively, insulators may not mediate homologue pairing *per se*, rather prior homologue pairing is a prerequisite for allowing insulators to mediate transvection. We favour the latter model: although we did not directly assay pairing, we have numerous instances where addition of extra copies of the GI has a detrimental effect on transvection (**Figures 6 and 7**). This result would be hard to reconcile with a model where insulators promote pairing, as we would expect pairing, and thus transvection, to locally increase as more and more insulators are added. Consistent with the view that homologue pairing precedes transvection is the fact that screens designed to identify somatic homologue pairing factors did not reveal any of the numerous insulator binding proteins (Bateman *et al.* 2012b; Joyce *et al.* 2012). Moreover, a recent study imaged two homologously inserted transgenes in live embryos and found the same frequency of colocalization (pairing) whether an insulator was included or not (Lim *et al.* 2018) – yet transvection between these genes required an insulator on both homologues, in agreement with our results. Another recent study mapped DNA elements mediating pairing and transvection from the *ss* locus: the two activities were found to map on two different fragments (Viets *et al.* 2018).

If insulators do not mediate homolog pairing, their role could be to enable the productive interaction of enhancers with certain promoters. Several of our observations support such a more active role: (1) In order to promote *trans*-interaction between *e8* and either *pH* or *p7*, the WI requires the presence of *another enhancer* (*e7*) nearby. (2) Transvection supported by single 5’ GIs is orientation-dependent. Having both homologues in the FOR orientation is much more effective than both homologues bearing a GI in the REV orientation (**Figure 6**); if insulator-insulator interactions were the only parameter influencing transvection, having congruently disposed insulators in both homologues would most likely produce an identical result, whether the configuration were FOR/FOR or REV/REV. (3) When two GIS are present in each homologue, they exhibit a strong bias for “vertical” trans GI/GI interactions (**Figure 7**). The fact that this bias can be alleviated by promoter mutations is consistent with more direct insulator-promoter communication. (4) A forward GI exhibits a strong promoter preference: it transvects the *3xP3* enhancer only to *pH* and not to *p7* or *p8* in a certain transgene combination (**Figure S9**).

Recent data agree with such a more direct role of insulators in enhancer-promoter communication. A genome-wide chromatin occupancy analysis for more than 15 insulator binding proteins showed that a large proportion of their binding sites is near a promoter or an enhancer (Cubeñas-Potts *et al.* 2016). Direct contacts between insulators and nearby enhancers and promoters has been detected in transgenes *via* 3C (Kyrchanova *et al.* 2013). Live imaging of two loci separated (in *cis*) by >100kb has shown that homotypic insulators promote proximity between these loci, but they do it much more effectively in the presence of a promoter in the one locus that gets activated by enhancers on the other (Chen *et al.* 2018). Finally, studies replacing specific insulator elements in the Bithorax Complex with other insulators, strongly support the ability of the resident insulators, like Fab-7 and Fab-8, to interact with neighboring enhancers (the *iab* elements) to bring them in the proximity of the *AbdB* promoter (Kyrchanova *et al.* 2016; Postika *et al.* 2018). How insulators select which enhancers to pair with which promoters is an important question that still remains to be elucidated.

Why has the necessity for insulators been overlooked in some of the studies on transvection? Most probably because the fly genome and common transgenesis cloning vectors are rich in insulators. For example, in one study (Mellert and Truman 2012) all transgenes used contained mini-*white* and its associated WI, which we have shown is capable of mediating some enhancer action in *trans*, consistently transvection was observed only with a subset of enhancers in that study. Two other studies (Bateman *et al.* 2012a; Blick *et al.* 2016) used recombinase-mediated cassette exchange system which allows for transgene integration without vector sequences, however, both of the landing sites used (JB53F and JB37B) are in close proximity (200 bp and 1 kb) to endogenous insulators bound by CTCF (Negre *et al.* 2010). In our experiments we never observed transvection in the absence of an insulator in the transgenic construct, even though we tested five different *attP* landing sites. Out of these five *attP* sites, the closest endogenous insulator to the regulatory elements present in our constructs is 6 kb away in the case of *VK2*, while in the case of the rest of the loci (*VK13, VK37, VK40* and *attP40*) this distance is no less than 20 kb, taking into consideration the vector sequences of the integrated transgenes (Negre *et al.* 2010). This suggests, not only a necessity of insulator elements for transvection, but also that they need to be closely linked to the transvecting enhancer and promoter; which agrees with the emerging genome-wide association of insulators with enhancers and promoters (Bartkuhn *et al.* 2009; Cubeñas-Potts *et al.* 2016). As a corollary, one should be careful when planning to use heterozygous transgene combinations in the same landing site. If one wishes to minimize transvection, one should preferably not include GI or any other insulator in the transgenes; at the expense of losing shielding from position effects. If shielding is desired, GIs can be used in only one of the transgenes. If shielding of both transgenes is desired, GIs can be placed in different orientations and as far as possible from the transgenes’ enhancers and promoters. If, on the other hand, one wishes to promote transgene transvection, one should place GIs in the forward orientation directly upstream of the transgenes’ cis-regulatory elements.

The association between insulators and transcriptional *cis*-regulatory elements (enhancers and promoters) is not a peculiarity of *Drosophila*; it has been reported also for vertebrates (Guo *et al.* 2015; reviewed in Hnisz *et al.* 2016). On the other hand, *Drosophila* (dipterans in general) seem to be unique in establishing somatic homologue pairing early in development and maintaining it throughout life (Abed *et al.* 2018; Erceg *et al.* 2018). Transvection could be an epiphenomenon of these two biological processes: insulator interactions with enhancers/promoters and homologue pairing. This would explain why it is more often encountered in *Drosophila*, but is only sporadic in mammals (Apte and Meller 2012; Stratigi *et al.* 2015). Does transvection also serve a role in regulating transcriptional output and accordingly could it be positively selected in evolution? Some studies have suggested that it increases transcription from the two alleles, or that it coordinates their transcriptional on/off decisions (Goldsborough and Kornberg 1996; Johnston *et al.* 2014). How widespread this effect is across the genome and whether it contributes to organism adaptation to fluctuations in environmental conditions or response to stressful stimuli is not known. At the least, transvection would ensure robustness of gene expression levels in the face of genetic variation, specifically heterozygosity for mutations in promoters or enhancers.

**Table 1.**
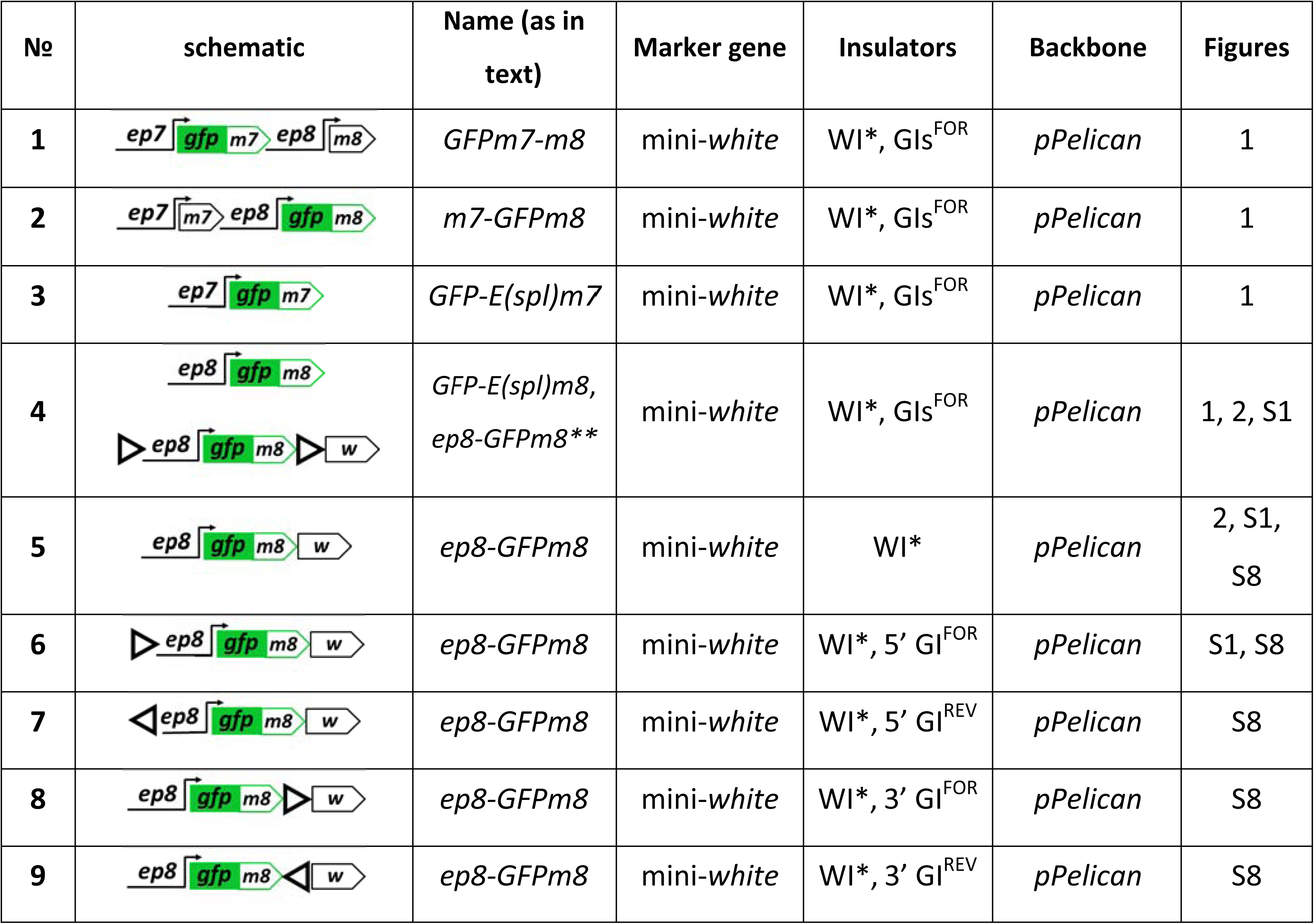
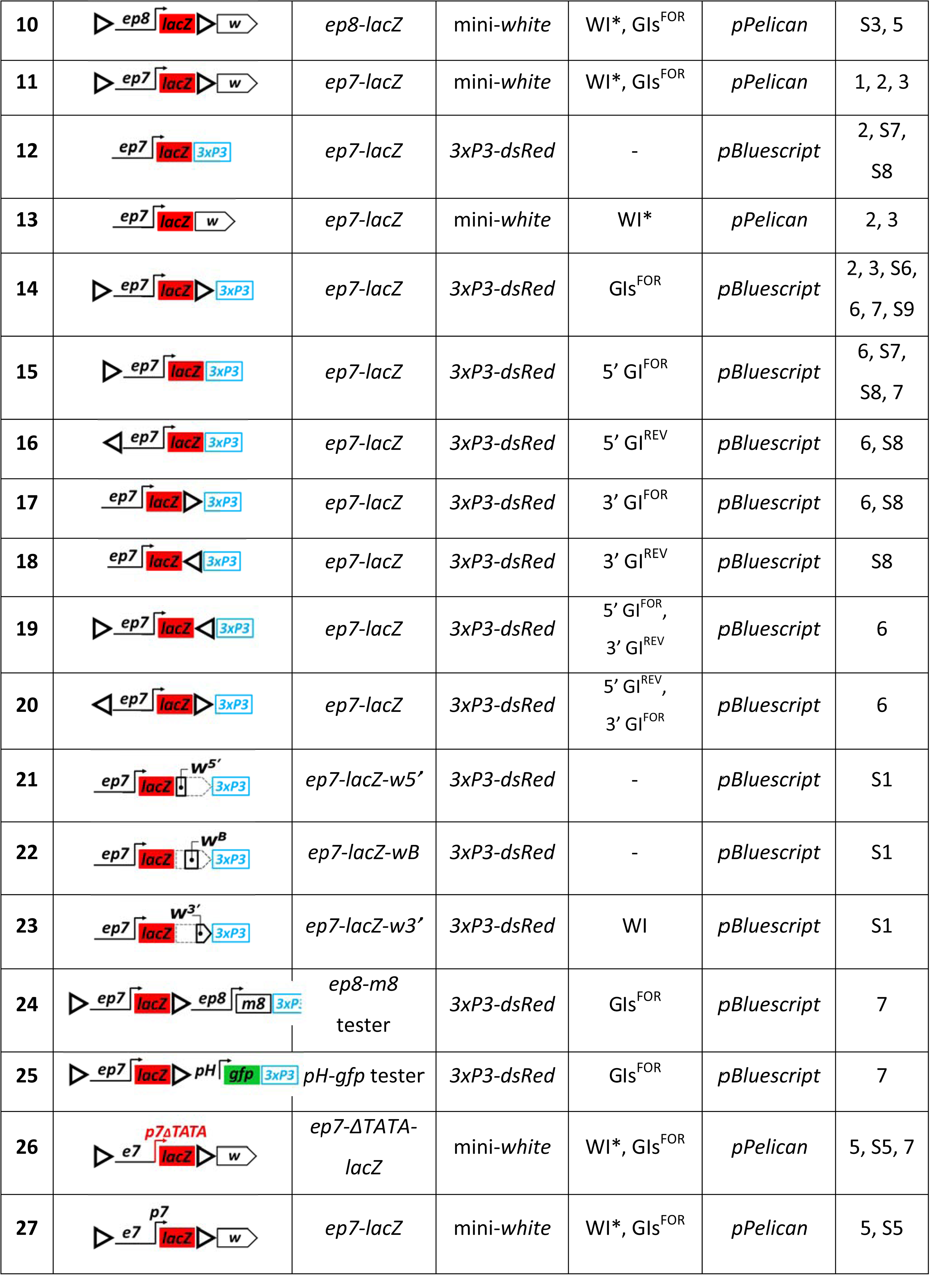
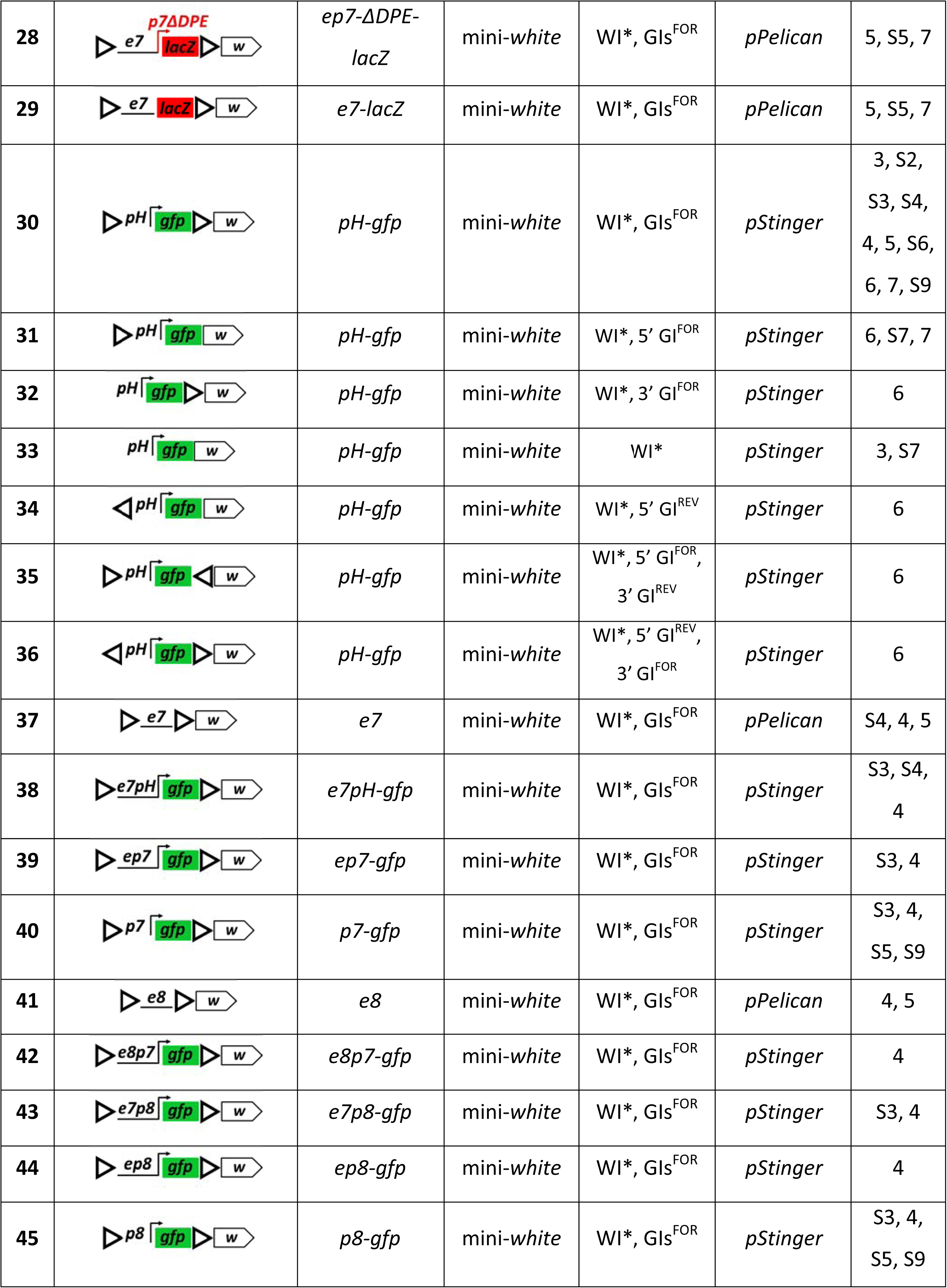
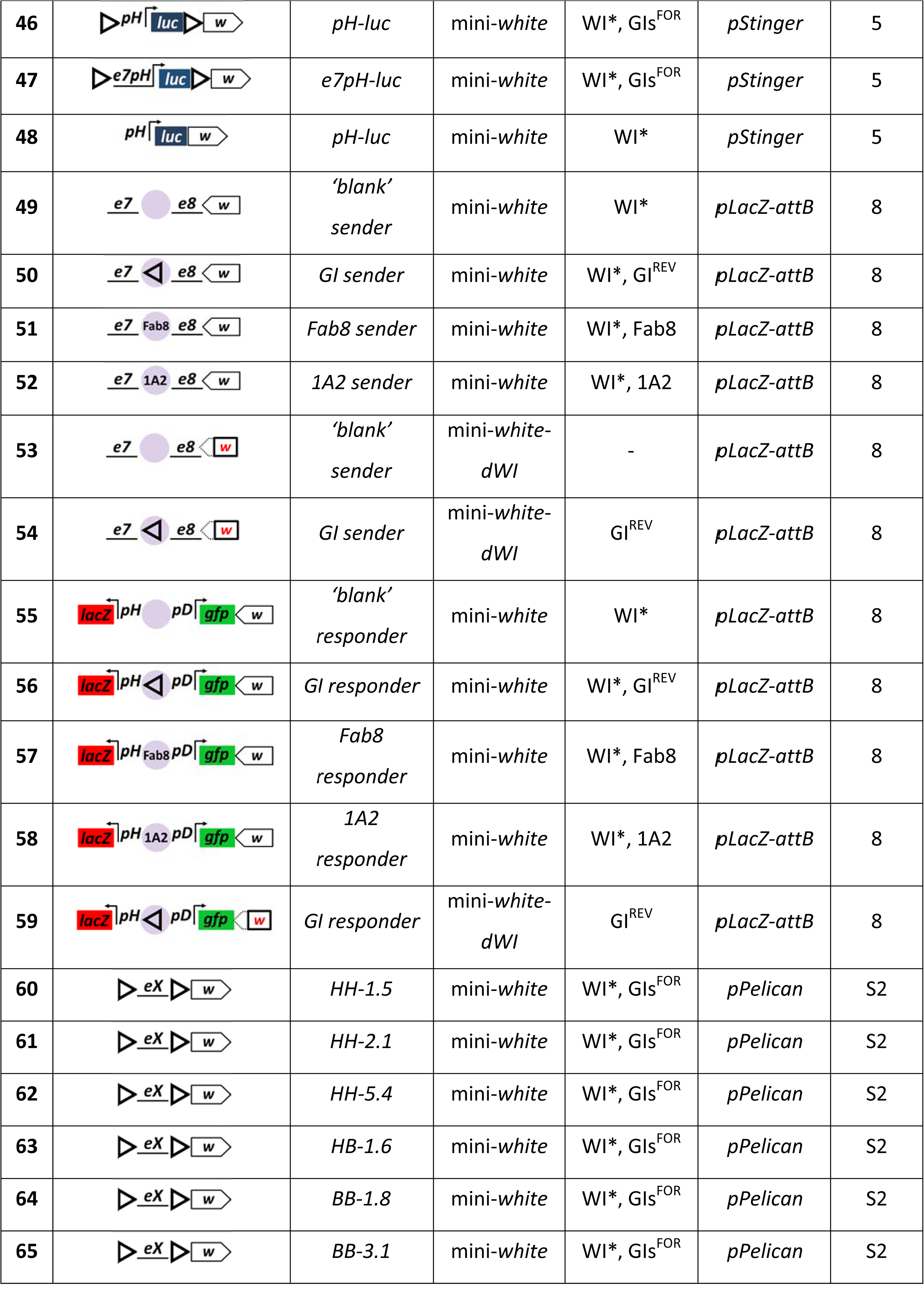
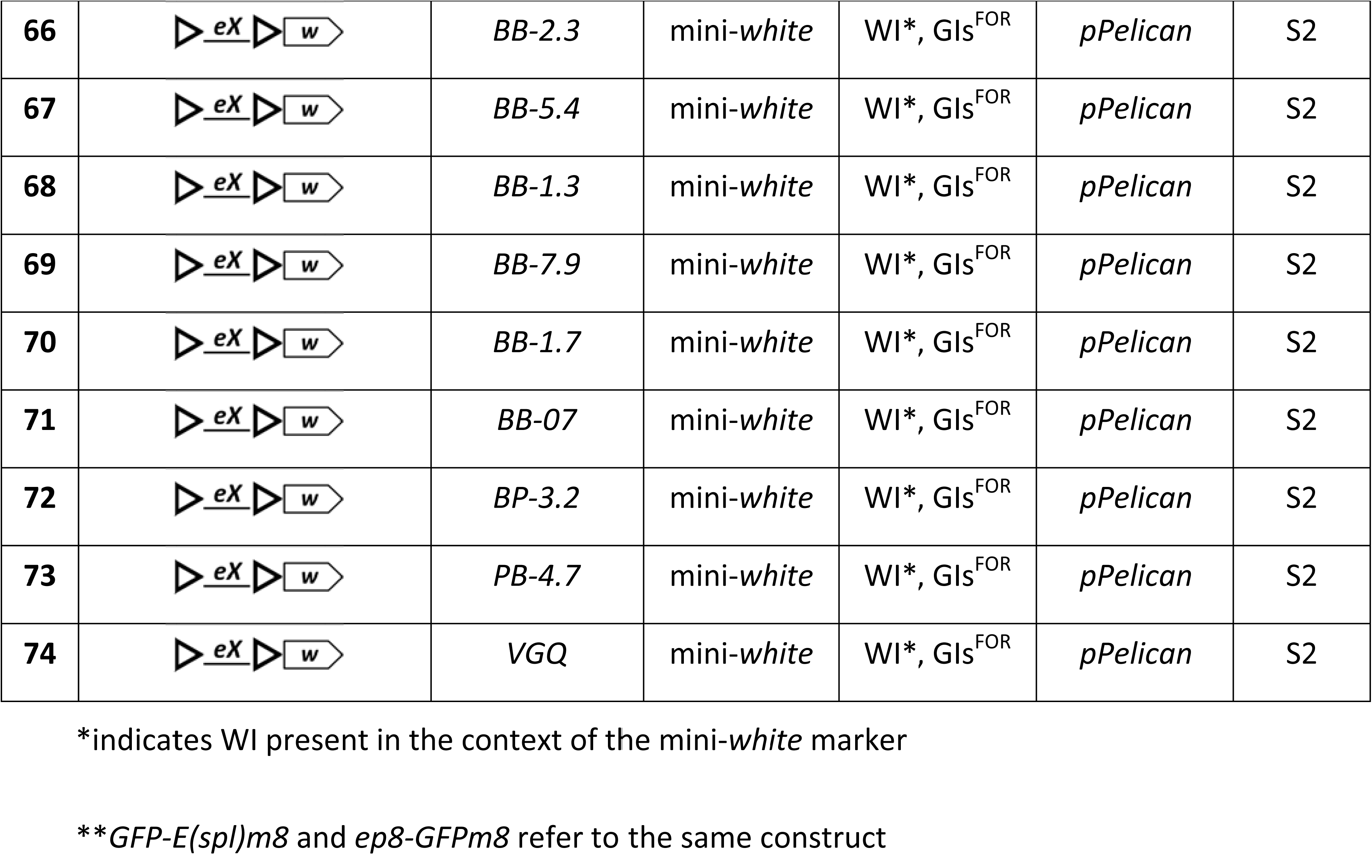
List of constructs used in the study

**Table 2.**
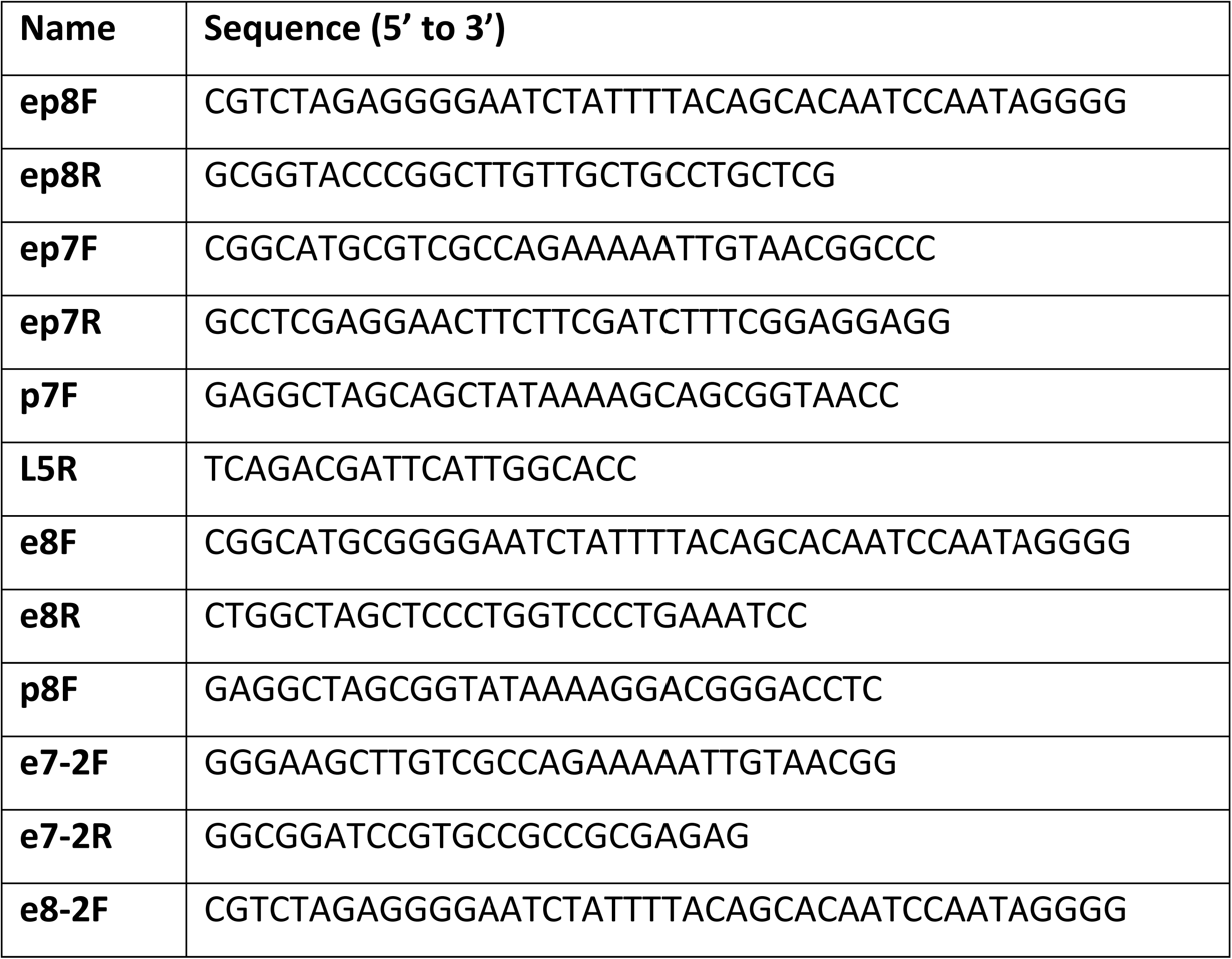
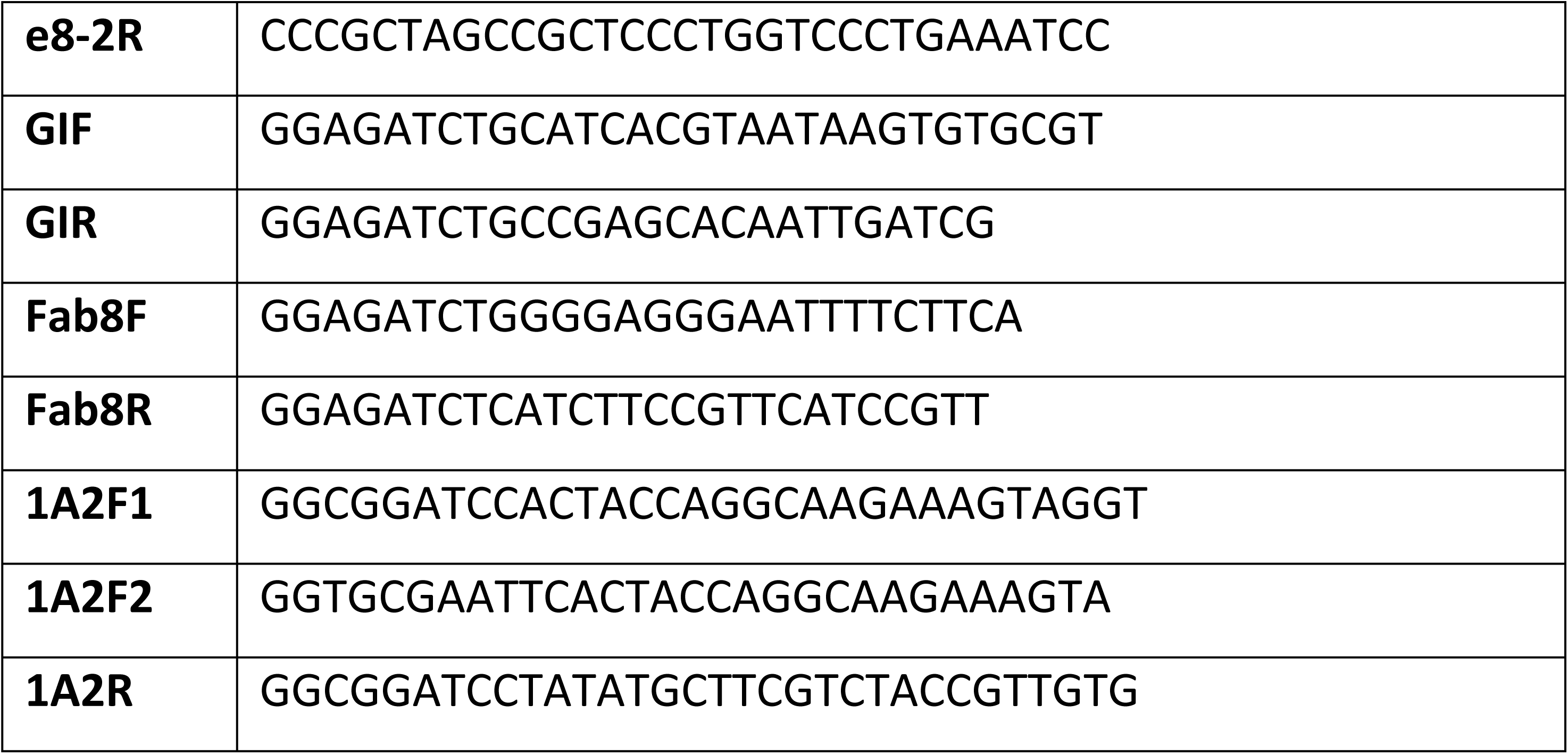
List of primers

## Acknowledgments

We thank Michalis Averof for *pMinos*{*3xP3-dsRed*}, Maria Monastirioti for *pGL3-hsp70-luc*, Chrysoula Pitsouli and Norbert Perrimon for providing *attP2* and *attP40* flies; Marina Gkantia, Giorgos Samantsidis, Alexis Molfetas, Babis Galouzis and Vasilis Ntasis for their help in generating plasmid constructs and isolating transgenic flies. Many thanks to Dr Charalampos Spilianakis for critical reading of the manuscript. This work was supported by the EU Marie Curie EST program No 7295/FAMED, ARISTEIA grants (No 4436 and 1967) from the General Secretariat for Research and Technology of Greece (co-funded by the EU European Social Fund), the Fondation Santé and intramural IMBB support.

## SUPPLEMENTARY MATERIALS AND METHODS

**The role of insulators in transgene transvection in *Drosophila***

**Piwko et al**

### Plasmid constructs

Here we present the general strategy used for plasmid construction. Detailed information regarding each construct and maps are available upon request. **Table 1** is the list of all constructs used in this study together with their general characteristics, provides numeric (**№**) identification for each construct and associates them with the Figures in the Results section. **Table 2** provides information of the primers used in plasmid construction.

To obtain our constructs we have used four different backbones of publicly available plasmids: *egfp*-containing *pStinger* and *lacZ*-containing *pPelican* (Barolo *et al.* 2004), *pBluescript* (*SK*(*+*), Stratagene) and *pLacZ-attB* (Bischof *et al.* 2007). *pStinger, pPelican* and *pBluescript* were modified by cloning *attB* sequence from *pTA-attB* plasmid (Groth *et al.* 2004) in the same orientation in each plasmid, immedietaly upstream to the multi-cloning site (MCS; in case of *pStinger*/*pPelican* it is upstream of the 5’ GI).

The 7 kb genomic fragment encompassing *E*(*spl*)*m7* and *E*(*spl*)*m8*, together with their upstream (enhancers and promoters) and downstream (3’ UTRs) regulatory regions, was cloned with *XhoI* and *ClaI* restriction enzymes from the R3012 cosmid clone encompassing part of *E*(*spl*) locus (Delidakis and Artavanis-Tsakonas 1992) into *pPelican-attB*. Due to strong post-transcriptional repression of both of these genes (Lai *et al.* 2005) we have replaced *E*(*spl*)*m7* and *E*(*spl*)*m8* 3’ UTRs with the SV40 (derived from pGL3 vector, Promega) and the *Adh* (derived from the *Ract-HAdh* vector, Swevers *et al.* 1996) poly A terminators, respectively. Subsequently, unique restriction sites were introduced before the start codons of both genes by mutagenesis to allow cloning of EGFP (derived from *pCRE-d2EGFP* vector, Clonetech) into the ORF of *E*(*spl*)*m7* in one construct (*GFPm7-m8*, contruct **№ 1** in Table 1) and of *E*(*spl*)*m8* in another (*m7-GFPm8*, construct **№ 2**).

The ‘short genomic’ constructs (*GFPm7* and *GFPm8*) were generated by the excision of the regulatory and coding portions of the untagged genes from the ‘long genomic’ constructs (i.e., *GFPm7* (construct **№ 3**) was generated from *GFPm7-m8* by a deletion of the downstream sequences to the *m7*’s (SV40) 3’ UTR, and *GFPm8* (construct **№ 4**) was generated from *m7-GFPm8* by a deletion of the *m7*’s 3’ UTR and its upstream sequences). The resulting constructs contained 2.1 kb sequence upstream of the start codon of *GFPm7* (which we denote as *ep7*) and 1.3 kb sequence upstream of the start codon of *GFPm8* (*ep8*).

The GI-less *ep8-GFPm8* construct (**№ 5**) was generated by cloning of the *ep8-GFPm8* module (together with its *Adh 3’ UTR*) from *GFPm8* into *pPelican-attB* bearing deletion of both GIs and LacZ. This construct was subsequently used to generate four single-GI versions of *ep8-GFPm8* by cloning the 5’ GI from *pPelican* into 5’ (upstream of *ep8*) or 3’ (downstream of *Adh 3’ UTR*) positions in forward or reverse orientation (constructs **№ 6-9**).

The *ep8-lacZ pPelican*-based construct (**№ 10**) was made by cloning PCR-amplified *ep8* (primers **ep8F** and **ep8R**) into the MCS of *pPelican*.

The *ep7-lacZ pPelican*-based construct (**№ 11**) was made by cloning PCR-amplified *ep7* (primers **ep7F** and **ep7R**) into the MCS of *pPelican*. The *pBluescript* GI-less version of *ep7-lacZ* (**№ 12**) was made by (*1*) cloning the entire *ep7-lacZ* sequence of *pPelican*-based *ep7-lacZ* (together with its *SV40 3’ UTR*) into *pBluescriptSK*(*+*) in between *KpnI* and *SpeI* restriction sites; (*2*) and subsequent cloning of the DNA fragment containing *3xP3-dsRed* and *attB* seqences from *pMinos*{*3xP3-dsRed*} vector (Berghammer *et al.* 1999; this vector was beforehand modified by us by inserting *attB* sequence in the vicinity of *3xP3-dsRed*) into *KpnI* site of the *pBluescript* such that *attB* site is in between the *3xP3-dsRed* and *ep7*. The GI-less version of the *ep7-lacZ* construct with mini-*white* marker (**№ 13**) was made by the replacement of the *3xP3-dsRed* in the latter construct with *pPelican*’s mini-*white*. The *pBluescript* GI-less version of *ep7-lacZ-3xP3-dsRed* (**№ 12**) construct was used as a basis to generate (*1*) GI-containg *ep7-lacZ* constructs by cloning 5’ GI from *pPelican* immediately upstream of *ep7* or/and downstream of *SV40 3’ UTR* in a reverse or forward orientation (constructs **№ 14-20**); (*2*) the three *ep7-lacZ* constructs containing different parts of mini-*white* gene cloned into *SpeI* site (immediately downstream of *SV40 3’ UTR*): the 0.24 kb *AfeI*/*EcoRI* fragment encompassing the 5’/promoter region of mini-*white* (*ep7-lacZ-w5’*, construct **№ 21**), the 2.4 kb *AflII*/*EcoRV* fragment encompassing mini-*white*’s gene body (*ep7-lacZ-wB*, construct **№ 22**) and the 0.9 kb *EcoRV*/*BsrGI* 3’ part of the mini-*white* (*ep7-lacZ-w3’*, construct **№ 23**). The *pBluescript ep7-lacZ* construct with two GIs in forward orientation (**№ 14**) was used to generate two ‘tester’ constructs by cloning *ep8-m8-Adh 3’ UTR* module from the EGFP-untagged version of the *GFPm8* construct (*ep8-m8* tester, **№ 24**) and *pH-gfp*-*SV40 3’ UTR* module from *pHStinger* (*pH-gfp* tester, **№ 25**) immediately downstream of the 3’ GI.

The *ep7-ΔTATA-lacZ* construct (**№ 26**) containing deletion of 20 nt encompassing TATA box was generated by introducing an *EcoRV* restriction site by site-directed mutagenisis upstream of the TATA box of *p7* in *pPelican*-based *ep7-lacZ* construct (**№ 11**) and subsequent excision of the seqeunce between *EcoRV* and *BstEII*. The intermediate *ep7-lacZ* construct with introduced *EcoRV* (but without *EcoRV*/*BstEII* deletion; **№ 27**) produced LacZ expression pattern and levels indistinguishable to that of the *ep7-lacZ* without *EcoRV* and was used as a control transgene (to *ep7-ΔTATA-lacZ* and *ep7-ΔDPE-lacZ*) containing wild-type *p7*. The *ep7-ΔDPE-lacZ* (**№ 28**) construct was generated from *pPelican*-based *ep7-lacZ* by excision of 83 nt containing INR and DPE motifs in between *BstEII* and *StuI* restriction sites. The *e7-lacZ* (promoterless construct, **№ 29**) was generated by an excision of sequence between *EcoRV* and *StuI* from the *ep7-lacZ* construct with the introduced *EcoRV* site (**№ 27**).

All *pH-gfp* constructs (**№ 30**-**36**) were made based on *pHStinger* (a version of *pStinger* containing *pH* fused to *egfp*; Barolo *et al.* 2000). The *pH-gfp* with two GIs in forward orientation (**№ 30**) was made by inserting *attB* sequence into *pHStinger*. Subsequent deletions of the 3’ GI, 5’ GI and both GIs from this construct resulted in generation of constructs **№ 31, 32** and **33**, respectively. The 5’ GI^REV^ *pH-gfp* (**№ 34**) was made by cloning GI in reverse orientation into GI-less *pH-gfp*. The 5’ GI^FOR^ 3’ GI^REV^ *pH-gfp* (**№ 35**) was made by cloning GI in a reverse orientation in place of the 3’ GI^FOR^ of GIs^FOR^ *pH-gfp* (**№ 30**). The 5’ GI^REV^ 3’ GI^FOR^ *pH-gfp* (**№ 36**) was made by cloning GI in a reverse orientation in place of the 5’ GI^FOR^ of the GIs^FOR^ *pH-gfp* (**№ 30**).

The 2 kb *BglII*-*EcoRV e7*-containing fragment derived from the *ep7-lacZ* construct bearing *EcoRV* site introduced upstream to TATA box of *p7* (**№ 27**) was used to generate (*1*) the *e7* construct (**№ 37**) by replacing *lacZ* in the *pPelican*-attB and (*2*) the *e7pH-gfp* (**№ 38**) by cloning it upstream to *pH* of the GIs^FOR^ *pH-gfp* (**№ 30**). The *ep7-gfp* construct (**№ 39**) was made by PCR-amplifying the *p7* promoter from *ep7-lacZ* (**№ 11**) with **p7F** and **p7R** primers and ligating the *NheI/HeaIII*-digested PCR product into *NheI* and *StuI* sites in the *e7pH-gfp* construct (**№ 38**) such that *p7* sequence replaces the sequence of *pH*. The *p7-gfp* (**№ 40**) was based on the *ep7-gfp* (**№ 39**) by excision of *e7* with *KpnI*. The sequence of *e8* was PCR-amplified with **e8F** and **e8R** primers, its product was cut with *AvrII* and *NheI* and ligated to *XbaI* and *NheI* sites (replacing *e7*) in the *e7* construct (**№ 37**) to generate *e8* construct (**№ 41**). The sequence of *e8* (as a *MfeI*-*NheI* fragment) was cloned from *e8* construct (**№ 41**) upstream of *p7* (*EcoRI*/*NheI*) in the *p7-gfp* construct (**№ 40**) to generate *e8p7-gfp* (**№ 42**). The *e7p8-gfp* construct (**№ 43**) was made by PCR-amplifying the *p8* promoter from *ep8-lacZ* (**№ 10**) with **p8F** and **L5R** primers and ligating the *NheI/HeaIII*-digested PCR product into *NheI/StuI* sites in the *e7pH-gfp* construct (**№ 38**) such that *p8* sequence replaces the sequence of *pH*. The *e8* sequence was cut out from the *e8* construct (**№ 41**) with *MfeI* and *NheI* restriction enzymes and ligated to *EcoRI* and *NheI* sites (such that the *e8* sequence replaces the sequence of *e7*) in the *e7p8-gfp* construct (**№ 43**) to generate *ep8-gfp* construct (**№ 44**). The *p8-gfp* construct (**№ 45**) was generated by excision of the *e7* sequence with *NaeI* and *NheI* restriction enzymes from the *e7p8-gfp* construct (**№ 43**).

The luc constructs (**№ 46**-**48**) were generated by replacing *pH-gfp* module with the restriction fragment containing *pH-luc* module (derived from the *pGL3-hsp70-luc* construct, gift from M. Monastirioti, IMBB) in the GIs^FOR^ *pH-gfp* (**№ 30**), GIs^FOR^ *e7pH-gfp* (**№ 38**) and GI-less *pH-gfp* (**№ 33**) constructs – resulting in generation of GIs^FOR^ *pH-luc* (**№ 46**), GIs^FOR^ *e7pH-luc* (**№ 47**) and GI-less *pH-luc* (**№ 48**), respectively.

The *‘blank’ sender* construct (**№ 49**) was made by cloning (*1*) the *HindIII/BamHI*-cut PCR-amplified product of *e7* (primers **e7-2F** and **e7-2R**) into *pLacZ-attB* construct (replacing *lacZ* sequence) and (*2*) subsequent cloning of the *NheI/BglII*-cut PCR amplicon of *e8* (primers **e8-2F** and **e8-2R**). The *‘blank’ sender* construct (**№ 49**) was used to generate GI, Fab8 and 1A2 sender constructs (**№ 50, 51** and **52**, respectively) by cloning the PCR-amplified sequences of GI, Fab8 or 1A2 into the *BglII* site located in between *e7* and *e8*. The 400 bp GI sequence was amplified on the *pPelican* template with **GIF** and **GIR** primers and contains full length GI. The 540 bp Fab8 sequence was PCR-amplified from *Drosophila* genome using **Fab8F** and **Fab8R** primers, which relates to genomic location 3R: 16918976..16919515 (*Dm6*), and contains the F8^254^ sequence and part of F8^469^ sequence (as defined in Kyrchanova *et al.* 2008). The 454 bp 1A2 sequence was PCR-amplified from *Drosophila* genome using primers **1A2F** and **1A2R**, and relates to the exact same region defined as 1A2 insulator in Kyrchanova *et al.* 2008a. The WI-less sender constructs, **№ 53** and **54**, were generated from the *‘blank’ sender* (**№ 49**) and the *GI sender* (**№ 50**) constructs, respectively, by excision of the 341 bp WI-containing fragment between *BsrGI* and *BstBI* sites.

The *‘blank’ responder* construct (**№ 55**) was made by (*1*) replacing the *pH* in the *pHStinger* with the the *DSCP* promoter (*pD*) derived from *pBPGUw* plasmid (Pfeiffer *et al.* 2008) and (*2*) ligating the resulting *pD-gfp-SV40 3’ UTR* module in between the *BglII* and *NheI* sites of *pLacZ-attB*, which already contains the *pH-lacZ-SV40* module (Bischof *et al.* 2007). Subsequently, this construct was used to generate *GI* and *Fab8 responder* constructs (**№ 56** and **57**, respectively) by cloning into the *BglII* site the sequences of GI and Fab8 PCR-amplified with the same sets of primers (i.e., **GIF**/**GIR** and **Fab8F**/**Fab8R**) and in the same orientation as it was for the corresponding *sender* constructs. The 1A2 sequence was amplified with the **1A2F2** and **1A2R** primers and ligated into *EcoRI/BglII* sites of the *‘blank’ responder* construct (**№ 55**) resulting in generation of the *1A2 responder* construct (**№ 58**). The WI-less *GI responder* (**№ 59**) was generated by a deletion of the 502 bp WI-containing *BstBI/NsiI* fragment from the *GI responder* construct (**№ 56**).

The *HH-1.5, HH-2.1, HH-5.4, HB-1.6, BB-1.8, BB-3.1* and *BB-2.3* constructs (**№ 60-66**) were generated by cloning 1.5 kb-, 2.1 kb-and 5.4 kb-*HindIII-HindIII*, 1.6 kb-*HindIII-BglII*, 1.8 kb-, 3.1 kb- and 2.3 kb-*BglII-BglII* fragments, respectively, derived from the *R3007* cosmid (Delidakis and Artavanis-Tsakonas 1992) into the MCS of *lacZ*-deficient *pPelican-attB* (not listted).

The *BB-5.4, BB-1.3, BB-7.9, BB-1.7, BB-07, BP-3.2* and *PB-4.7* constructs (**№ 67-73**) were generated by cloning 5.4 kb-, 1.3 kb-, 7.9 kb-, 1.7 kb- and 0.7 kb-*BglII-BglII*, 3.2 kb- and 4.7 kb-*PstI-BglII* fragments, respectively, derived from the *R3012* cosmid (Delidakis and Artavanis-Tsakonas 1992) into the MCS of *lacZ*-deficient *pPelican-attB*.

The 0.8 kb sequence of Vestigial Quadrant Enhancer (*vgQ*, Kim *et al.* 1996) was derived from a *pBluescript-vgQ* vector and cloned into the MCS of *lacZ*-defic ent *pPelican-attB* to generate the *VGQ* construct (**№ 74**).

---

**Figure S1.**
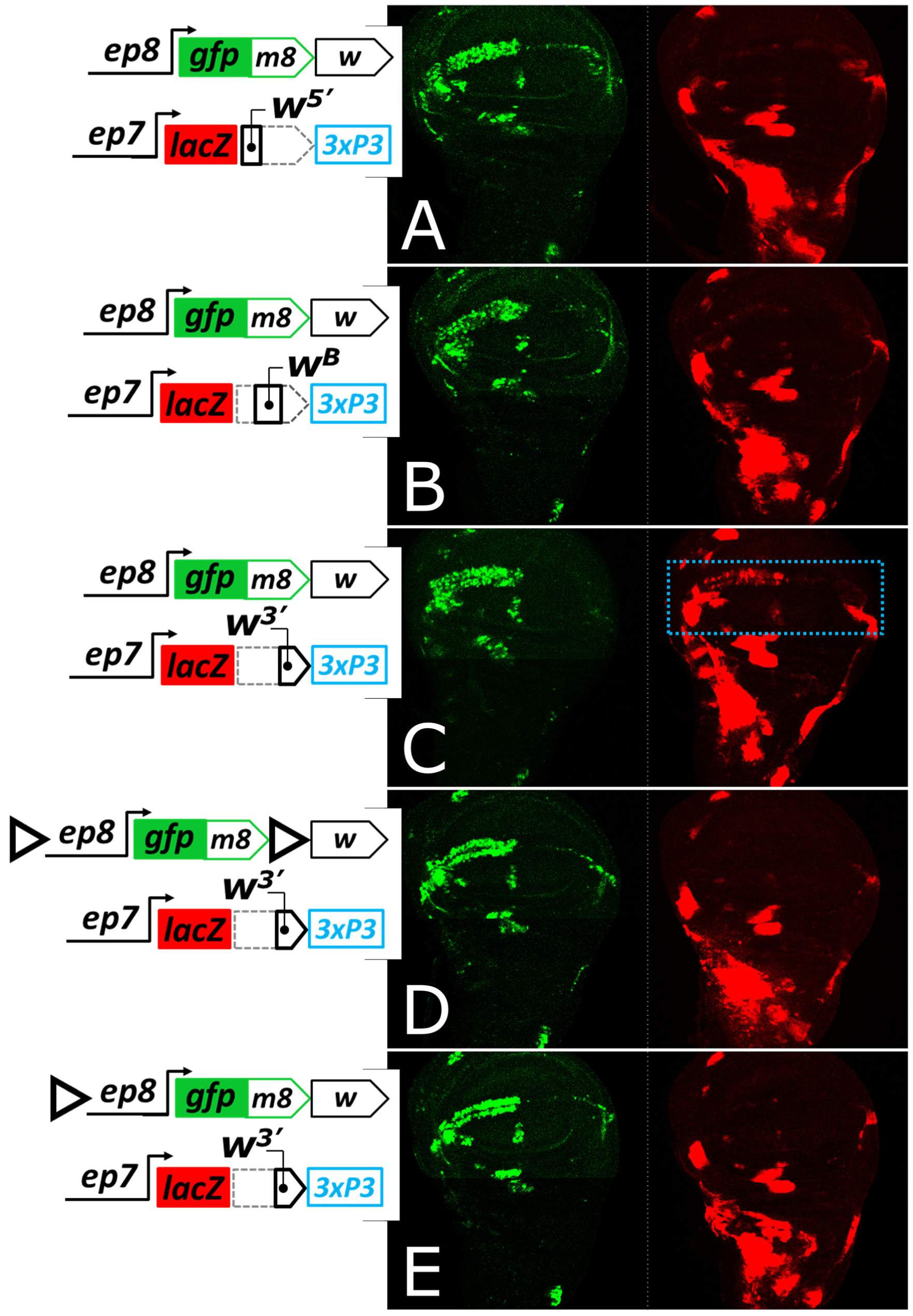
*White*-mediated uni-directional transvection relies on mini-*white*-contained Wari insulator (WI). Three fragments of mini-*white*, cloned in the *ep7-lacZ*(*-3xP3*) construct, were tested at the *attP40* locus in transvection assay with a *ep8-EGFPm8* transgene containing full-sized mini-*white* (**A**-**C**). Neither promoter region of mini-*white* (*w*^*5’*^, 0.24 kb, **A**), nor its ‘gene body’ (*w*^*B*^, 2.4 kb, **B**) mediated transvection; whereas the 3’ part of mini-*white* (*w*^*3’*^, 0.9 kb, **C**) did exhibit uni-directional transvection in WM (blue dotted rectangle). This fragment is known to contain WI. Addition of one or two GIs in the *ep8* construct inhibited WI-mediated transvection (**D**-**E**).

**Figure S2.**
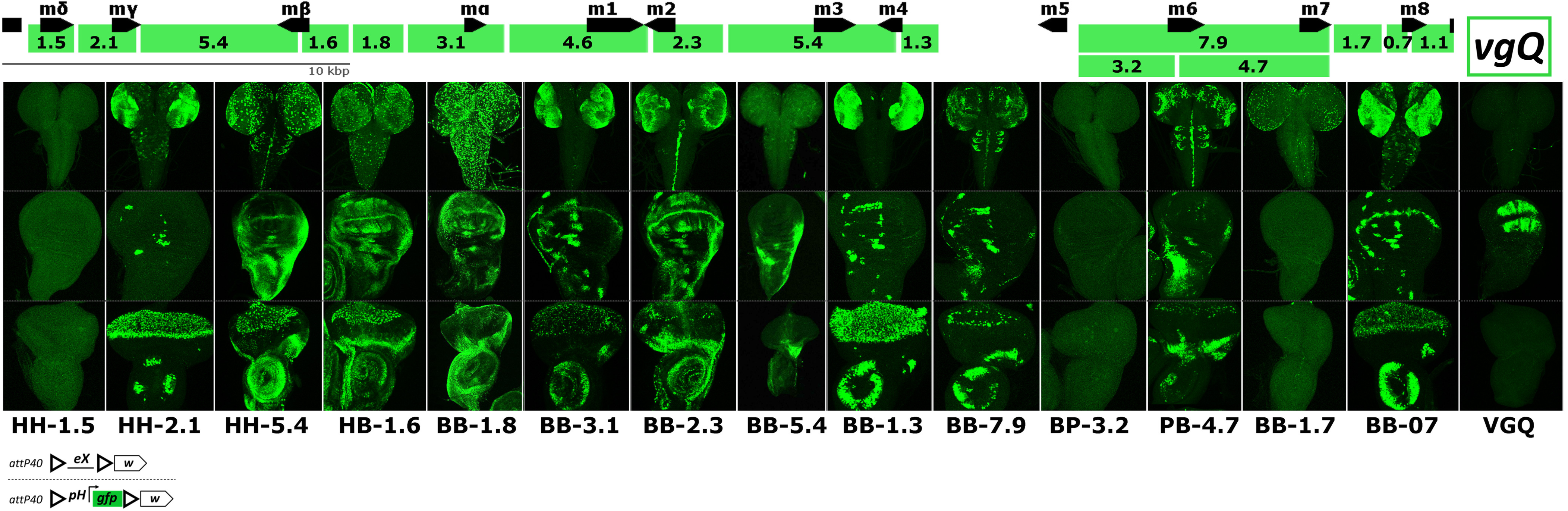
GI-mediated transvection is compatible with many developmental enhancers. A map of the *E*(*spl*) locus is shown on top and DNA fragments are marked as green rectangles with indicated length in kb. Each of these DNA fragments (’*eX*’ in the constructs’ scheme) was inserted into a transgene between GIs, placed in *trans* to GIs-flanked *pH-gfp* transgene in *attP40*. Heterozygotes were tested for activating *pH-gfp* expression in larval CNS (top), wing disk (middle) and eye-antennal disk (bottom row). Another fragment, unrelated to *E*(*spl*) locus, the vestigial quadrant enhancer (*vgQ*, last column) recapitulates its *cis* activity in *trans* (GFP expression only in the wing disk). We observe little or no activity from 3 *E*(*spl*) fragments in *trans*.

**Figure S3.**
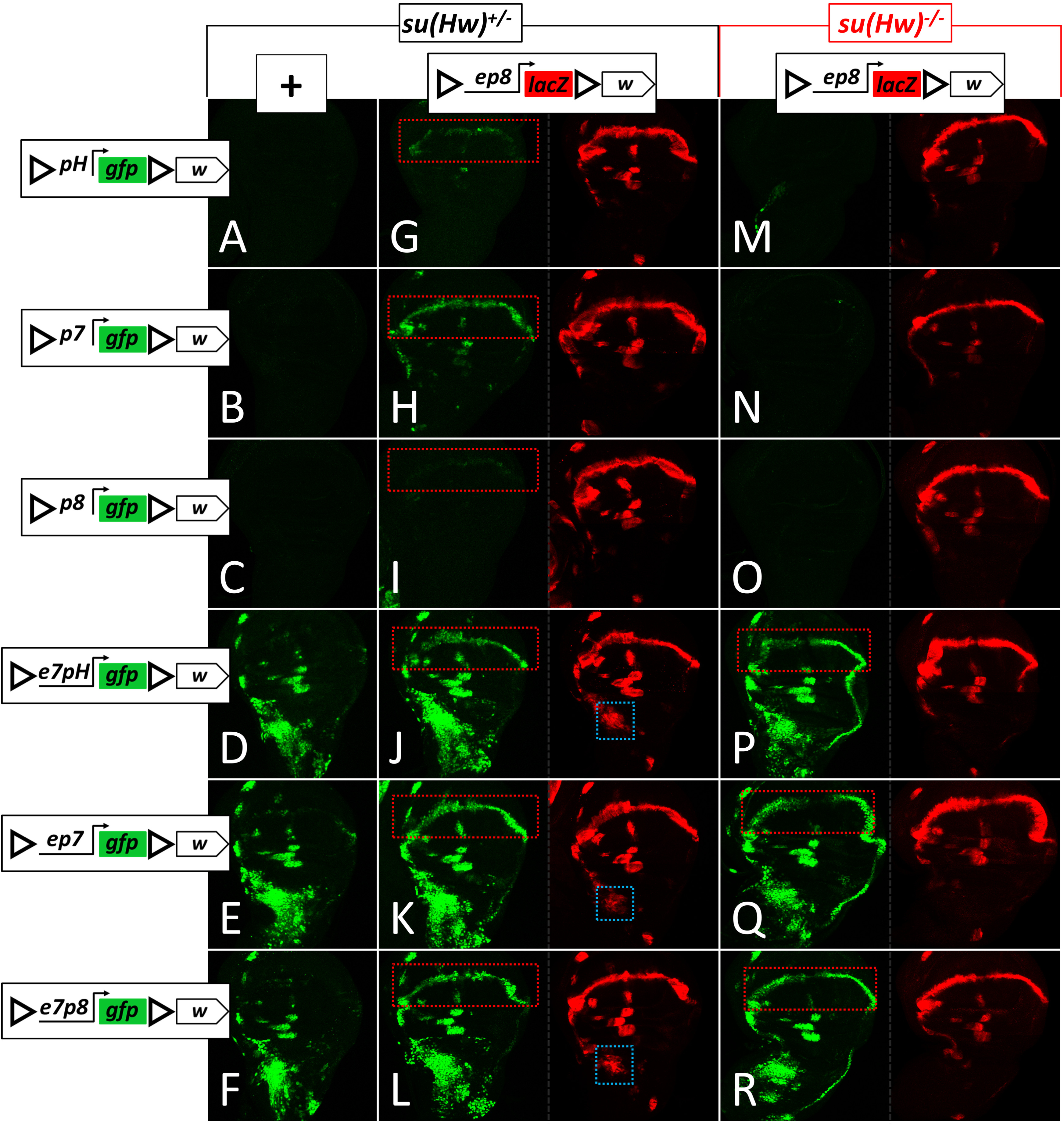
WI-mediated *trans*-activation by the *e8* enhancer requires the presence of the *e7* enhancer on the other homolog. All transgenes contain GIs and mini-*white* and are inserted in *attP40*. (**A**-**F**) Wing disks from animals hemizygous for the indicated GFP transgenes. When GFP is driven by the *pH* promoter (**A**), *E*(*spl*)*m7* (*p7*) promoter (**B**) and *E*(*spl*)*m8* (*p8*) promoter (**C**), no expression is observed; addition of the *e7* enhancer to any of these promoters (**D**-**F**) results in *e7*-specific expression pattern of the GFP; note the lack of WM expression. **G**-**L** show wing disks from animals heterozygous for the same *gfp* **A**-**F** transgenes and *ep8-lacZ.* **M**-**R** show the same transgene combinations in disks derived from *su*(*Hw*)^*-/-*^ mutants. Blue and red dotted rectangles highlight cells exhibiting *trans*-activity of *e7* (in AMPs) and *e8* (in WM), respectively. In the absence of *e7*, only GI-mediated transvection is detected (**G**,**H**,**I**), whereas in the presence of *e7* both GI- (**J**,**K**,**L**) and WI-mediated (**P**,**Q**,**R**) transvection takes place. WI-mediated transvection is unidirectional, as it is only observed in the WM (GFP) of **P, Q, R**, but not in their AMPs (LacZ); compare to bidirectional transvection in **J, K, L**.

**Figure S4.**
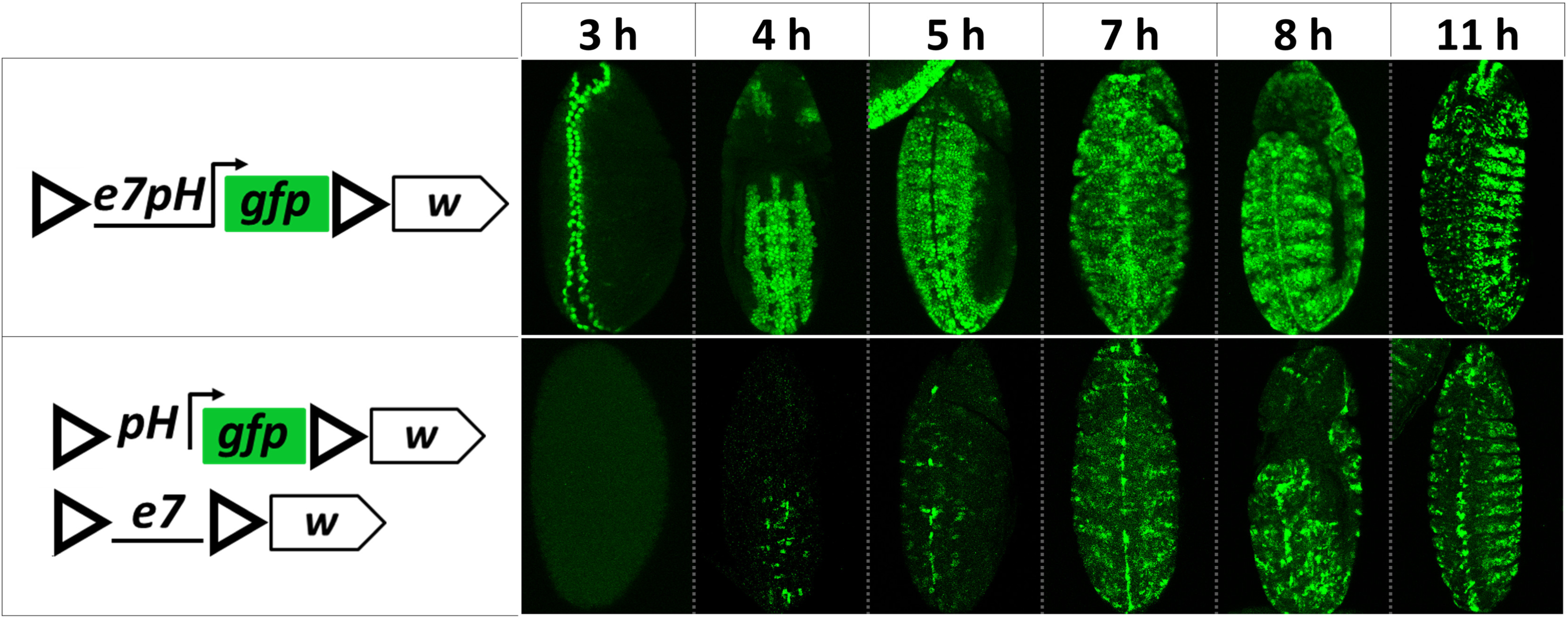
The onset of transvection is delayed in embryogenesis. All embryos are imaged ventrally with anterior to top. A *cis*-linked *e7pH* enhancer-promoter pair (top row) drives GFP expression along the ventral midline within 3 h after egg deposition (AED). At 4-5 h AED *e7pH* is broadly active in the ventral ectoderm, whereas *e7* and *pH* separated in *trans* show interaction only in small subset of these cells. From 7h AED onwards *e7* and *trans-pH* seem to interact in most cells where *e7*(*cis-*)*pH* is active. The enhancerless *ph-gfp* reporter shows no background expression in embryos as a hemizygote (data not shown).

**Figure S5.**
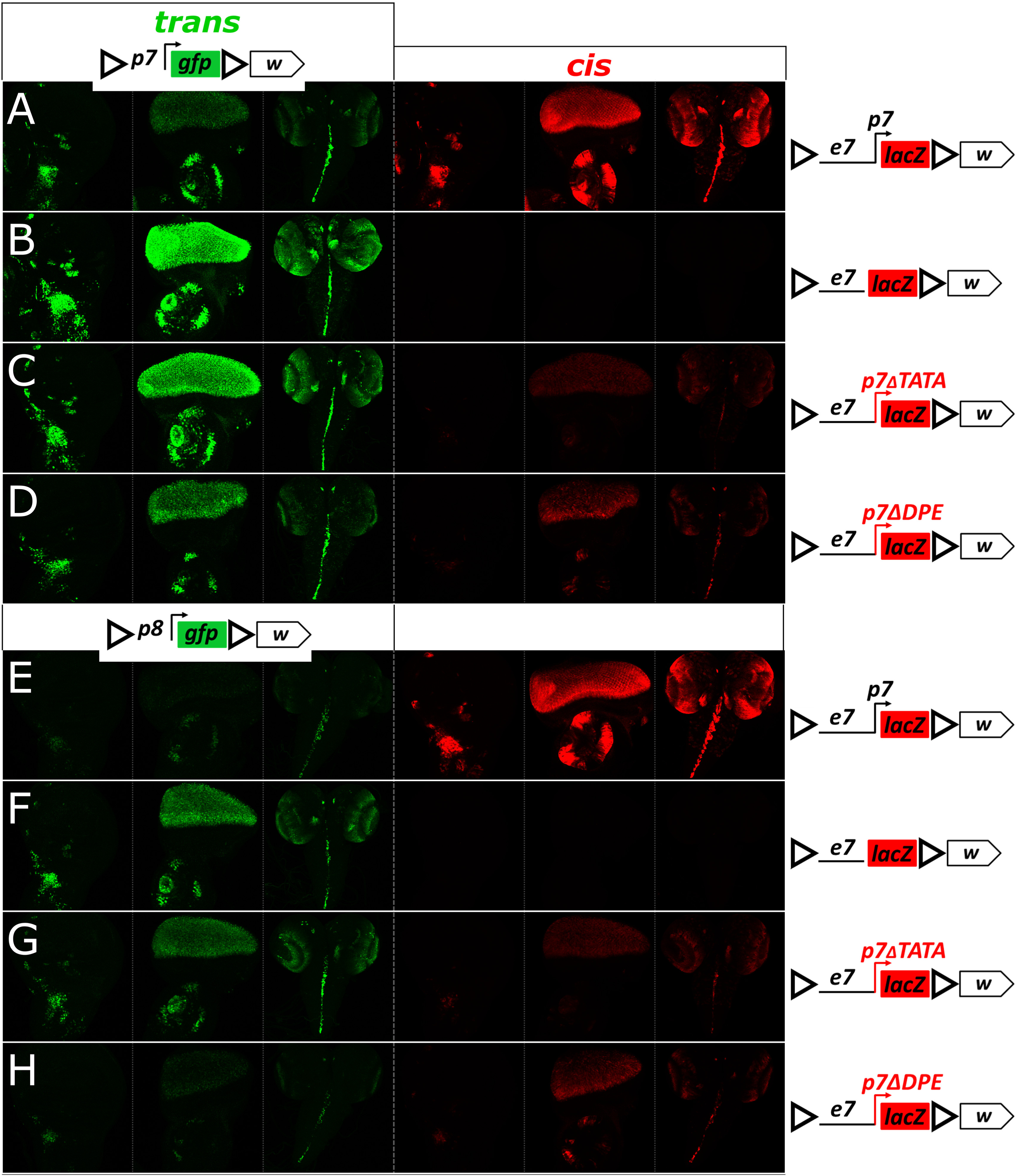
The effects of mutations in the *p7* promoter on the ability of a cis-linked *e7* to transvect are independent of the identity of the *trans* promoter. The *ep7-lacZ* (**A** and **E**) and its derivatives harboring deletions of *p7* (**B** and **F**), *ΔTATA* (**C** and **G**) and *ΔDPE* (**D** and **H**) are placed in *trans* to *p7-gfp-* (**A** - **D**) or *p8-gpf-*enhancerless constructs (**E** - **H**). For each genotype (each row) the third instar wing disk, eye disk and CNS are simultaneously examined for (*1*) GFP, relecting *trans* activity of the enhancer-containing transgenes on *pH-gfp* (green) and (*2*) β-galactosidase, reflecting *cis* activity the e7-linked promoter (*lacZ*, in red). Although, *p7, p8* and *pH* are promoters of different strength, their activity seems to be affected similarly by different mutations in *e7*-linked *trans*-*p7* (compare to the results obtained for *pH*, **Figure 5, D**-**G**). All transgenes contain GIs and mini-*white* and are inserted in *attP40*.

**Figure S6.**
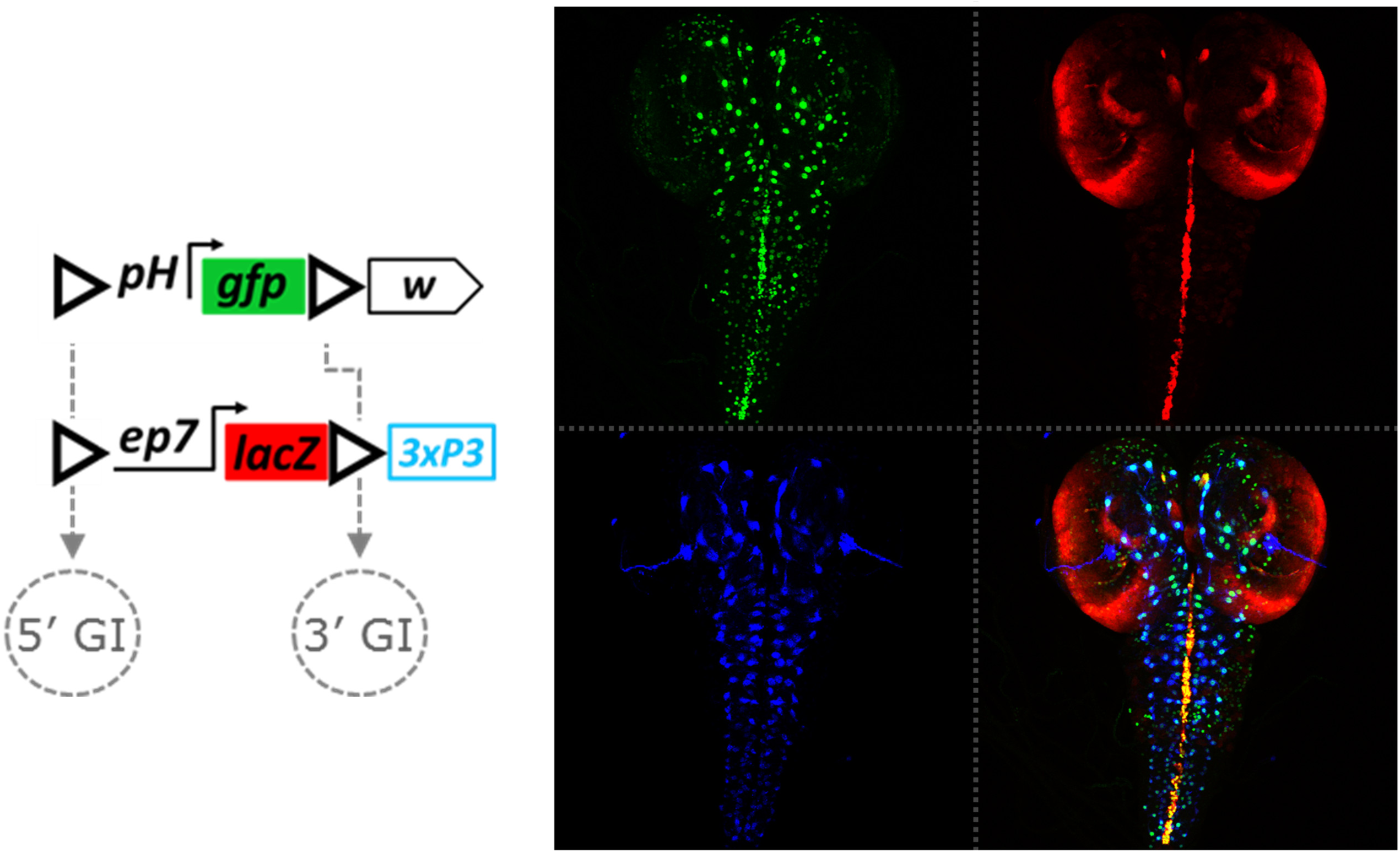
*pH* receives input from two enhancers in *trans, e7* and *3xP3*. Confocal z-projection of a third instar CNS from a heterozygote between *ep7-lacZ-3xP3* and *pH-gfp* dual-GIs^FOR^ transgenes in *attP40*. Top left panel shows *trans*-activated *pH-gfp* expression; note *e7*-specific GFP expression in the VNC midline, corresponding to the *cis*-activity of *e7* (driving expression of LacZ, in red, top right panel), and the ‘dotty’ *3xP3*-specific expression in central brain and VNC corresponding to glial cells with active *3xP3* enhancer (expressing DsRed, in blue, in *cis*). Merged image is shown in the bottom right panel.

**Figure S7.**
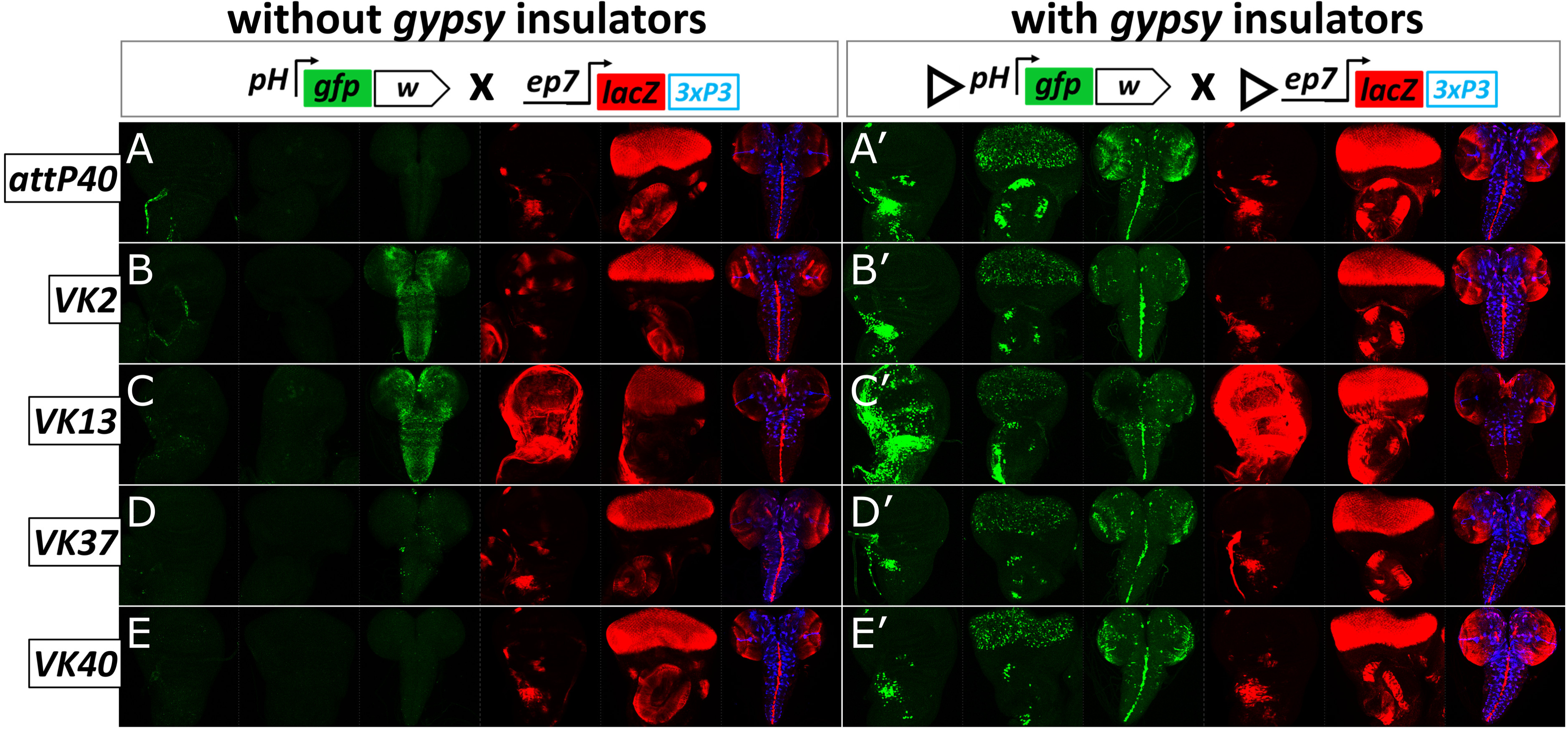
No transvection is observed in the absence of GIs at five different loci. The ability to interact in *trans* was tested between *ep7-lacZ* and *pH-gfp* transgenes inserted in five genomic loci (*attP40, VK2, VK13, VK37, VK40*) in two configurations of these transgenes: carrying no GIs (**A**-**E**) and containing one 5’ GI each (**A’**-**E’**). For each genotype three tissues are shown: wing disk, eye disk and CNS (left to right). Green is *pH*-driven GFP. Red is *ep7*-driven LacZ. Blue is *3xP3-*driven DsRed (CNS only). Note the activity of neighboring (trapped) enhancers in all loci except *VK40*, which is also seen in the hemizygous condition for these transgenes (not shown): (*1*) tracheal activity in the wing disk of the uninsulated *pH-gfp* in *attP40* and *VK2* and of the GI-insulated *pH-gfp* in *VK37* (**A, B** and **D’**), (*2*) ubiquitous activity in VNC and central brain of the uninsulated *pH* in *VK2* and *VK13* (**B** and **C**), (*3*) wing pouch activity in wing disk of the uninsulated *p7-lacZ* in *VK2* (**B**), (*4*) ubiquitous wing disk activity of uninsulated and GI-insulated *p7-lacZ*, and GI-insulated *pH* in *VK13* (**C** and **C’**). Note GFP expression in notum AMPs, retina and antenna and brain and VNC midline in **A’-E’.** All this e7-driven GFP expression is absent in **A-E**.

**Figure S8.**
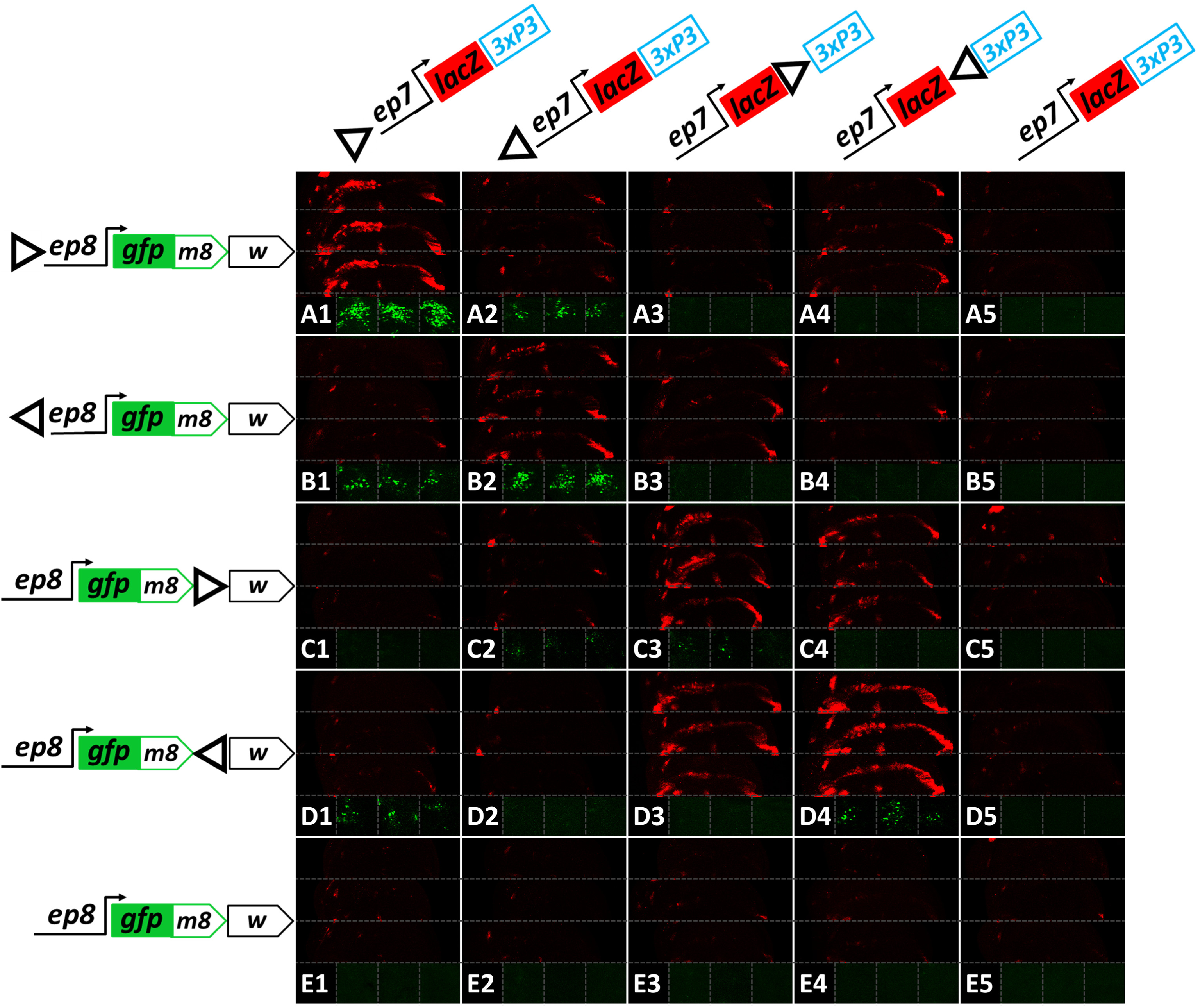
Less strict GI position and orientation requirements for interaction between *ep7* and *ep8*. (**A1**-**E5**) *ep8*- and *ep7*-containing single-GI transgenes in *attP40* are combined as shown. Each panel consists of six sub-panels containing confocal z-projections of the WM (only LacZ channel, in red) or the AMPs (only GFP channel, in green) from three different wing disks for each genotype. Both of these expressions result from transvection and are abrogated when one or both homologs lack a GI (row **E** or column **5**). Note, that unlike *ep7*→*pH* transvection (**Figure 6 H**), *ep8*→*ep7* transvection (LacZ in the WM) is also mediated by the 3’GIs (**C3, C4, D3, D4**). The 3’ GIs also mediate weak *ep7*→*ep8* transvection (GFP in AMPs), when congruently oriented (**C3, D4**). Note, also, that the absolute orientation of 5’ GIs is less important for *ep7*→*ep8* transvection (compare GFP in **A1, A2, B1, B2**) than for *ep7*→*pH* (**Figure 6I, L**).

**Figure S9.**
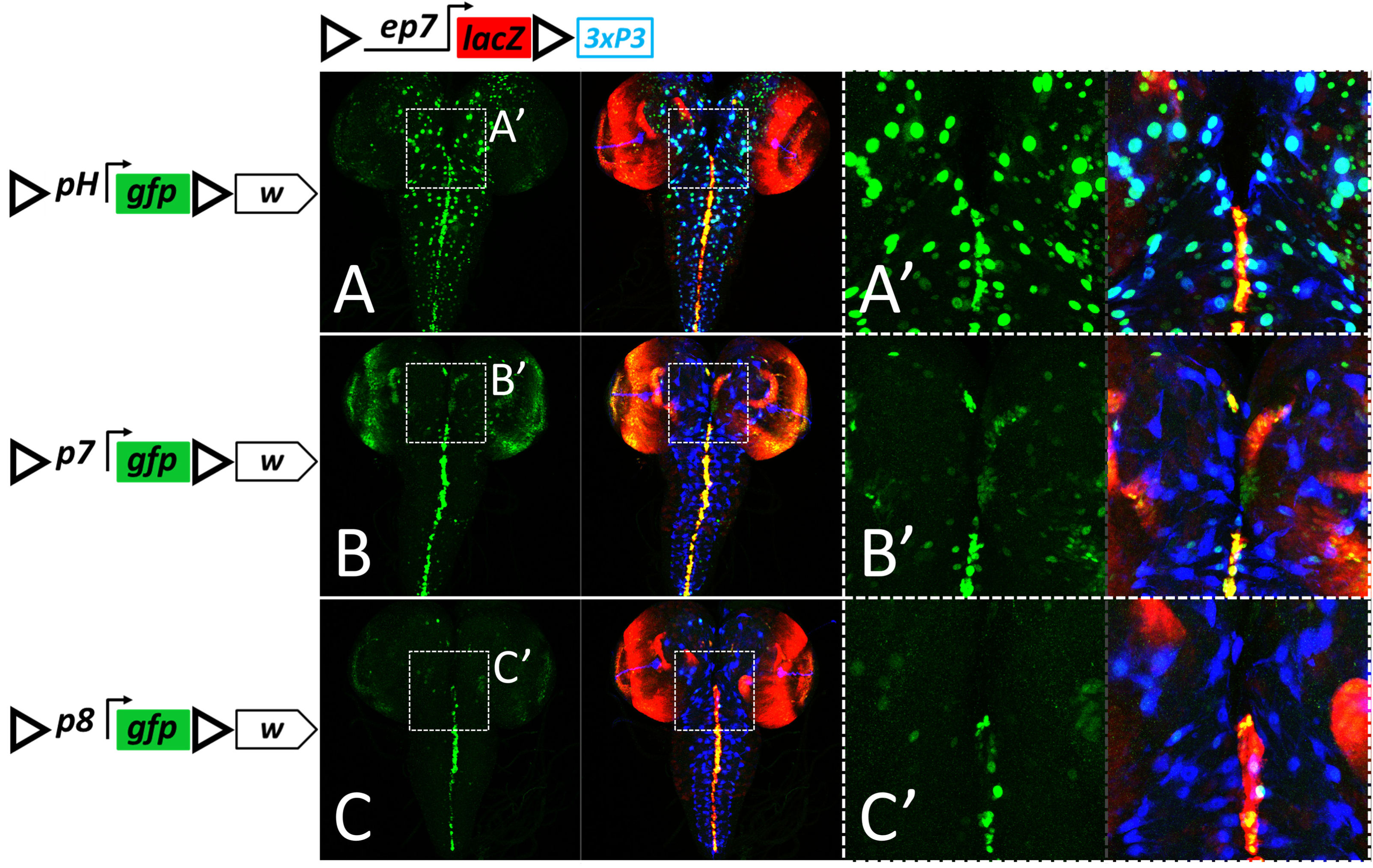
The *3xP3* enhancer has a stronger affinity for *pH* than for two other promoters. (**A**-**C**) Confocal z-projections of third instar CNSs showing GFP channel alone (green) and merged channels of GFP (green), LacZ (red) and DsRed (blue). (**A’**-**C’**) Enlarged regions of the central brains indicated by white rectangles in **A**-**C**. Out of the three enhancerless dual-GI^FOR^ *gfp* transgenes, each carrying a different promoter (**A** - *pH*, **B** - *p7*, **C** - *p8*), only the *pH*-driven reporter shows robust interaction with *3xP3* in the dual-GIs^FOR^ *ep7-lacZ-3xP3* transgene in *attP40*, giving a dotty glial GFP pattern. Notice weaker and sporadic GFP expression from *trans-3xP3* by *p7* (**B, B’**), even though *p7* is a stronger promoter than *pH*. In contrast, the “vertical” interaction of *e7* with the trans promoter (GFP in VNC midline and optic lobes) is stronger with *p7* (**B**) than with *p8* or *pH* (**A, C**).

